# How Myosin VI Traps its Off-State, is Activated and Dimerizes

**DOI:** 10.1101/2023.06.30.547185

**Authors:** Louise Canon, Carlos Kikuti, Vicente J. Planelles-Herrero, Tianming Lin, Franck Mayeux, Helena Sirkia, Young il Lee, Leila Heidsieck, Léonid Velikovsky, Amandine David, Xiaoyan Liu, Dihia Moussaoui, Emma Forest, Peter Höök, Karl J. Petersen, Aurélie Di Cicco, Julia Sires-Campos, Daniel Lévy, Cédric Delevoye, H. Lee Sweeney, Anne Houdusse

## Abstract

Myosin VI (Myo6) is the only minus-end directed nanomotor on actin, allowing it to uniquely contribute to numerous cellular functions. As for other nanomotors, proper functioning of Myo6 relies on precise spatio-temporal control of motor activity via a poorly defined off-state and interactions with partners. Our structural, functional, and cellular studies reveal key features of myosin regulation and indicate that not all partners can activate Myo6. TOM1 and Dab2 cannot bind the off-state while, GIPC1 binds Myo6, releases its auto-inhibition and triggers proximal dimerization. Myo6 partners thus differentially recruit Myo6. We solved a crystal structure of the proximal dimerization domain, and show that its disruption compromises endocytosis in HeLa cells, emphasizing the importance of Myo6 dimerization. Finally, we show that the L926Q deafness mutation disrupts Myo6 auto-inhibition and indirectly impairs proximal dimerization. Our study thus demonstrates the importance of partners in the control of Myo6 auto-inhibition, localization, and activation.

## Introduction

Myosin motor proteins generate force and/or movement from ATP hydrolysis when associated with actin filaments. Conformational changes in the motor as it progresses from ATP hydrolysis to release of inorganic phosphate and ADP on actin are amplified into large movements via a calmodulin (CaM) or light chain binding region referred to as the “Lever arm” (Fig. 1A). To control the functions of myosin motors in cells, the ATPase activity of the motor and its ability to interact with actin must be regulated both spatially and temporally. Thirteen different classes of myosin motors serve diverse cellular functions in mammalian cells^1^. The regulation of their motor activity is however poorly characterized. A general theme for the control of motor activity is the formation of intra-molecular interactions involving the C-terminal Tail region and the Motor domain of these motors. In the active form of the motor, the Tail region interacts with itself or cellular partners. The best understood is the case of the dimeric myosin II (Myo2) class^2^ and myosin V (Myo5) class^3^. In cardiac muscle, impairment in the stabilization of the myosin off-state leads to severe cardiomyopathies^4^. Whether lack of regulation of unconventional myosins can also lead to pathology has not been demonstrated.

**Figure 1.**
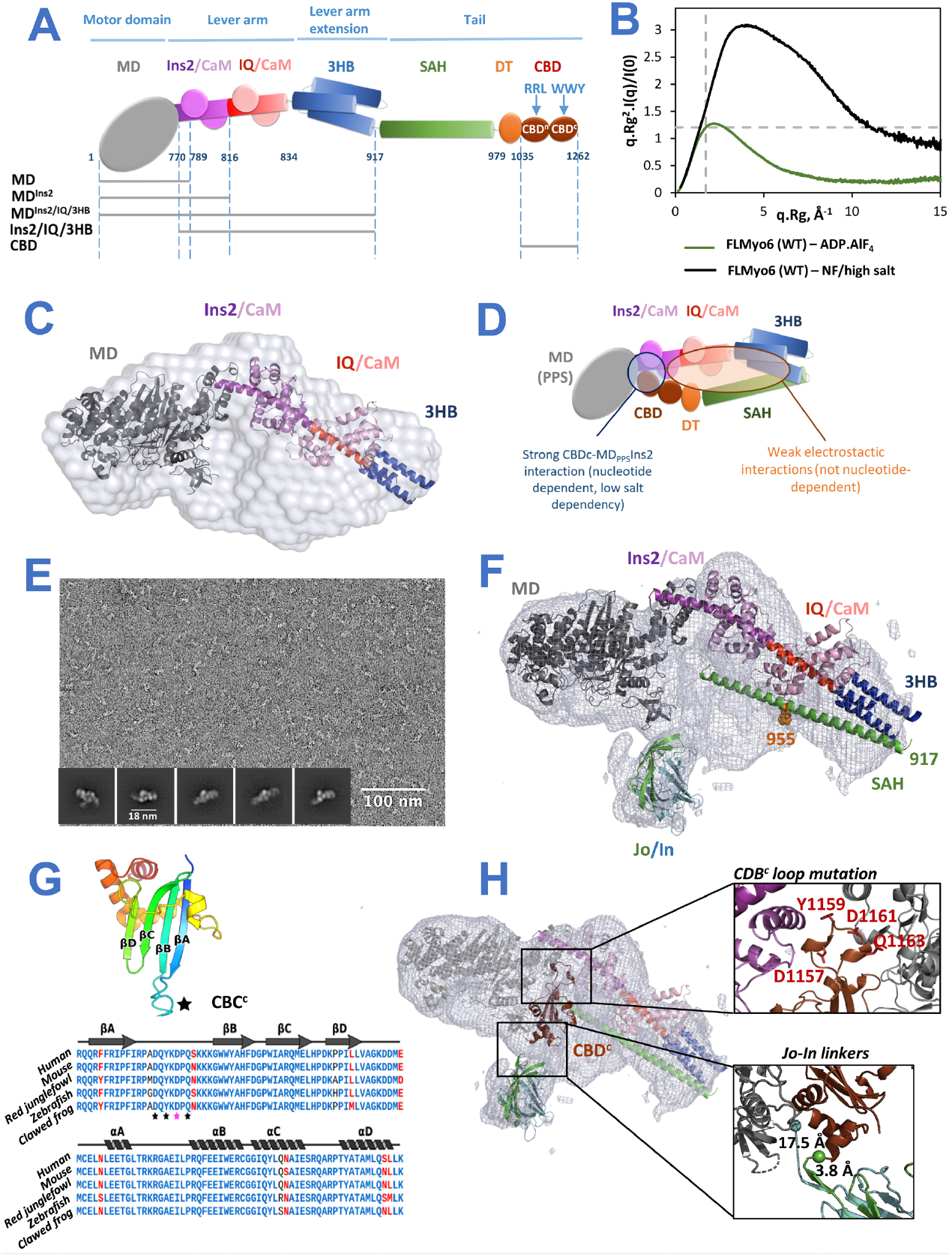
Importance of ADP.P_i_ for the compact, back-folded Myo6 conformation. **(A)** Schematic representation of FLMyo6 with the Motor domain (MD, grey), CaM binding sites (Ins2/IQ, purple/red), CaM (lilac/pink), 3-helix bundle (3HB, blue), single alpha helix (SAH, green), distal Tail (DT, orange) and CBD (brown). Residue numbers correspond to human Myo6, Uniprot entry Q9UM54-2. **(B)** Dimensionless Kratky plot representation from SEC-SAXS. FLMyo6 in the presence of ADP.AlF_4_ (green) results in a bell-shape spectrum with a maximum close to the intersection of the dashed lines (√3:1.104), typical of a globular protein. The spectrum for FLMyo6 in NF/high salt (black) suggests a much more elongated shape. See buffer compositions in Methods. **(C)** Representation of the ab initio SAXS envelope (light gray) with MD^Ins2-IQ-3HB^ docked. **(D)** Scheme representing the interactions stabilizing the Myo6 back-folded state. **(E)** Example of a negative staining micrograph of Jo-Myo6-In in ADP.VO_4_, with selected 2D classes. **(F)** EM density for Jo-Myo6-In (grey mesh) obtained by negative staining. Docking of Myo6 fragments and Jo-In (see Methods). **(G) (Top)** Crystal structure of the Myo6 C-terminus (CBD^c^) (PDB: 3H8D^14^). Star: highly conserved and exposed loop between the β_A_ and β_B_ strands. **(Bottom)** Alignment of Myo6 CBD^c^ domain (aa 1143 to 1262 in Q9UM54-2) from different species. Strictly conserved and similar residues are shown in blue and red, respectively. Stars: residues implicated in binding to the Myo6 Head (Table 1). **(H)** CBD^c^ (brown) added to the model pictured in (**F**) (see Methods).

Perhaps the most divergent form of regulation is emerging for Class VI (Myo6), VIIa (Myo7a), and X (Myo10) myosins, which all contain regions of extended stable single alpha helices (SAH). Indeed, while they are back-folded monomers in their inactive form^5–8^, these motors can self-associate to form active dimers upon activation^9–12^. How back-folding is stabilized is unknown and the nature of the dimerization following unfolding has only been elucidated for Myo10^10,12^, which forms an anti-parallel coiled-coil immediately following the SAH. The SAH thus extends the Lever arm in the case of Myo10^9,13^. Whether this is also the case for the dimeric Myo6 is debated and requires elucidation of its dimerization region^12,14–18^. The manner in which the motor is dimerized and the composition of its Lever arm greatly influence its function. A distinctive dimerization region in Myo10 allows the dimer to easily reach out for neighboring actin tracks and participate in filopodia formation^13^, unlike the vesicle transporter, Myo5, that makes multiple steps on a single actin track.

**Table 1.** Main contacts that stabilize the back-folded conformation Dissociation constant (K_D_) ± K_D_ confidence determined by microscale thermophoresis of Myo6 Head constructs against Myo6 Tail constructs (constructs schematized in Fig. 1A). Microscale thermophoresis profiles are presented in Sup Fig. 3.

| Head | Tail | Motor state | $K_D$ (nM) | n |
| --- | --- | --- | --- | --- |
| MD<sup>Ins2/IQ/3HB</sup> | YFP<sup>CBD</sup> | ADP.VO<sub>4</sub> | 144 ± 61 | 3 |
| MD<sup>Ins2</sup> | YFP<sup>CBD</sup> | ADP.VO<sub>4</sub> | 343 ± 197 | 3 |
| MD | YFP<sup>CBD</sup> | ADP.VO<sub>4</sub> | 3920 ± 1453 | 3 |
| Ins2/IQ/3HB | YFP<sup>CBD</sup> | - | 250 ± 86 | 4 |
| MD<sup>Ins2/IQ/3HB</sup> | YFP<sup>CBD</sup> | NF | 726 ± 480 | 2 |
| MD<sup>Ins2/IQ/3HB</sup> | YFP<sup>CBD<sub>D1157V.Y1159D.D1161R.Q1163V</sub></sup> | ADP.VO<sub>4</sub> | n.b. | 3 |

As a minus-end directed actin motor, Myo6 performs unique cellular roles (reviewed in^19^), including endocytic vesicle trafficking and maturation, stereocilia maintenance^20^ and melanosome maturation^21^, among many others. For these cellular functions, Myo6 must associate with different binding partners, such as Dab2, GIPC1 and TOM1 in distinct endosomal compartments^22–24^ and NDP52 in autophagy and in RNA Polymerase II transcription^25^. Initially characterized as a deafness gene^20^, Myo6 is also overexpressed in aggressive cancers^26,27^ and its depletion reduces cell migration and proliferation^26,27^.

Full-length Myo6 (FLMyo6) was characterized as a back-folded monomer *in vitro*^7,8^, which was confirmed to exist in cells by FLIM (Fluorescence lifetime imaging microscopy)^25^. TOM1 and Dab2 bind Myo6 through its WWY motif on the C-terminus part of its cargo-binding domain (CBD^c^; Fig. 1A) while GIPC1 and NDP52 bind the RRL motif on the CBD^n^ (Fig. 1A). FRET studies showing that these partners can unfold constructs lacking the Myo6 Motor domain (MD) led to the proposition that all partners could activate Myo6 upon binding^25,28,29^. However, whether the WWY and RRL motifs are both accessible in the FLMyo6 back-folded state is unknown. Detailed studies of Myo6 recruitment are required to investigate the role of partners in the spatio-temporal regulation of its cellular activity.

The configuration of the Myo6 active state, the nature of its Lever arm and its oligomeric state can make critical differences in the way the force produced by the motor is used^17,30,31^. In fact, Myo6 is well adapted to transport as well as to anchor, depending on the load it is working against, according to single molecule assays^32^. Although the capacity of partners to favor either monomeric, dimeric or oligomeric assemblies has been described^14,25,28,29,31,33–35^, the active configuration required to perform the different cellular roles of Myo6 is unknown. *In vitro* studies have identified a proximal dimerization region^12,18^, but its role for the cellular function of Myo6 is not established, nor is the structure of this region. Furthermore, it is not known whether the dimerization occurs following partner binding, and whether all partners lead to the same motor configuration, which ultimately will determine the nature of the effective Lever arm and mechanical performance of the motor.

A detailed description of the Myo6 off-state, a structural characterization of the proximal dimerization region, and the role of partners in Myo6 regulation are all essential to understand how Myo6 function is regulated in cells. Here we used structural and functional assays to thoroughly investigate these properties of Myo6. We demonstrate that not all partners can relieve Myo6 auto-inhibition since not all binding sites are accessible, and importantly we solved the structure of the proximal dimerization domain and demonstrate its validity.

## Results

### ADP.P_i_ bound to the motor strongly stabilizes the off-state conformation of Myo6

Previous biophysical characterizations of the Myo6 back-folded state identified contacts between the Myo6 Lever arm and CBD (Fig. 1A)^25,36^. However, a possible role of the Motor domain in back-folding remains to be clarified. *Size Exclusion Chromatography coupled with Multi-Angle Light Scattering* (SEC-MALS) and SEC *coupled with Small-Angle X-ray Scattering* (SEC-SAXS) experiments (Sup Fig. 1A-C; Fig. 1B) indicate that FLMyo6 adopts a compact conformation in the presence of ADP.P_i_ analogs (Radius of gyration (Rg) = 49.23 ± 0.92 Å) (Fig. 1B-C; Sup Fig. 1C) even at high salt concentration (~425 mM NaCl) (Sup Fig. 1A-B). In contrast, when no nucleotide is present (nucleotide-free (NF) condition), FLMyo6 shifts from a compact to an elongated conformation in a salt concentration-dependent manner (at high salt, Rg = 84.18 ± 4.33 Å, elution 1 mL earlier from SEC-MALS) (Fig. 1B; Sup Fig. 1A-C). Overall, high salt dependency of FLMyo6 opening, combined with the lack of salt dependency in presence of a nucleotide, suggests that the Lever arm and the Tail are held together *via* electrostatic interactions, while the interactions that keep the Tail back-folded on the Head require the Motor domain to be in a nucleotide-bound state of its cycle (Fig. 1D). At very low salt (10 mM KCl) and in the presence of actin, FLMyo6 consumes ATP ~10 fold slower than the Tail-less construct MD^Ins2^ (Fig. 1A; Sup Fig. 1D), indicating that the back-folded state is auto-inhibited.

### 3D reconstruction of the Myo6 off-state

To further characterize the Myo6 off-state, negative staining electron microscopy (EM) of FLMyo6 in ADP.VO4 (ADP.Pi analog) resulted in heterogeneous 2D classes, likely due to the intrinsic flexibility of the protein particles. Previous FLIM demonstrated that fusion of the N- and C-termini to fluorescent proteins is compatible with Myo6 back-folding^25^. Thus, we fused the N- and C-termini of Myo6 to two covalent bonding subdomains of *Streptococcus pneumoniae* pilus adhesin RrgA: Jo and In^37^ (the Jo-Myo6-In construct) to attempt to limit the inherent flexibility of the back-folded monomer.

To show that the fusion does not disrupt the Myo6 back-folding, we confirmed that the Jo-Myo6-In heavy chain could bind to two CaM using SDS-PAGE and that the Jo-Myo6-In behaved as a compact folded protein, even in NF/high salt condition using SEC-MALS and SEC-SAXS (Sup Fig. 2A-C). An actin-activated stopped-flow experiment revealed a low P_i_ release rate for Jo-Myo6-In compared to FLMyo6 wild-type (WT), indicating that the conformational changes required to release P_i_ were greatly slowed (Sup Fig. 2D). Finally, negative staining EM images of Jo-Myo6-In in ADP.VO_4_, low salt were collected (Fig. 1E). The 3D reconstruction of the Myo6 off-state at ~17 Å resolution (Fig. 1F, Sup Movie 1) is consistent in shape and dimensions with SAXS data of FLMyo6 (Fig. 1C, F).

**Figure 2.**
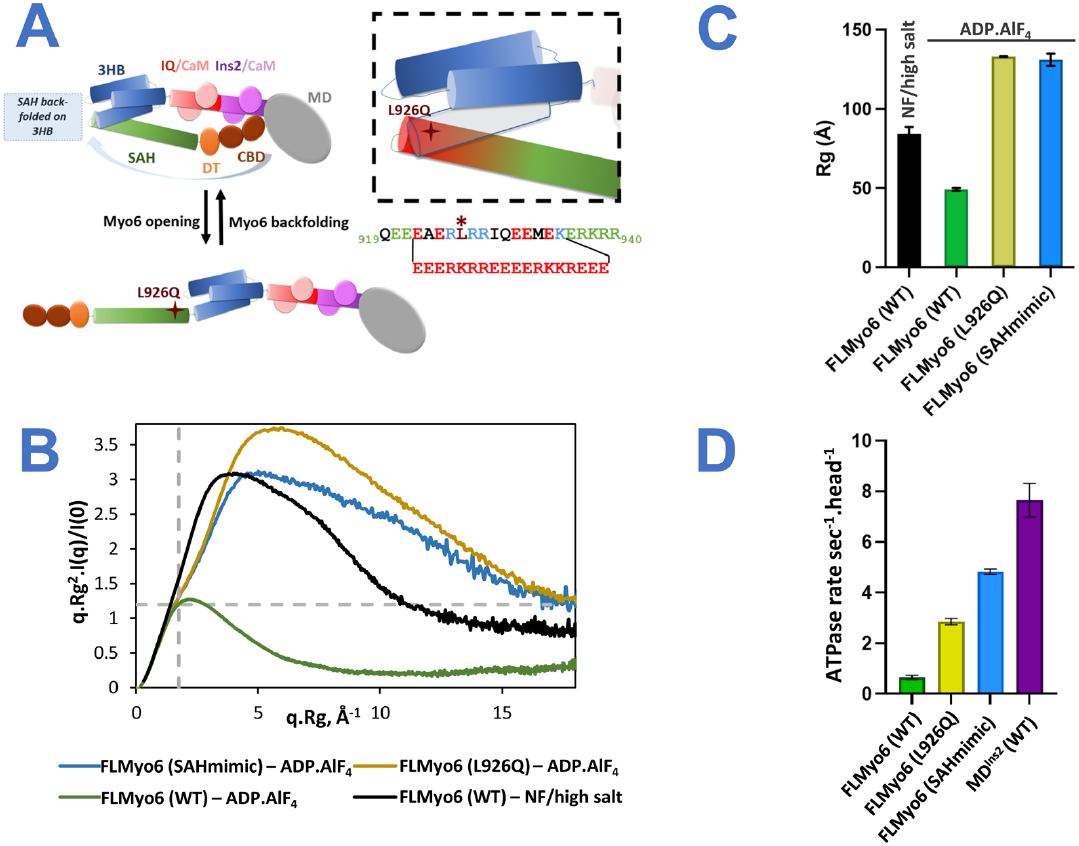
Role of the proximal Myo6 sequence in the stabilization of the Off-state. **(A)** Model of Myo6 opening/back-folding. Back-folding requires the SAH to fold back on the 3HB. The L926 residue (red cross) leads to deafness when mutated into Gln^38^. **(Insert)** Mutations of the apolar residues at the N-terminus of the SAH to convert it into a « perfect SAH» sequence (SAHmimic) turn Myo6 into a constitutive monomer^17^. **(B) (Top)** Dimensionless Kratky plot representation from SEC-SAXS. FLMyo6 in NF/high salt is pictured in black. In the presence of ADP.AlF_4_, FLMyo6 (L926Q) (yellow) and FLMyo6 (SAHmimic) (light blue) spectrums correspond to an elongated shape, as opposed to FLMyo6 WT (green). **(Bottom)** Rg of FLMyo6 WT, L926Q and SAHmimic determined by SEC-SAXS experiments in the presence of ADP.AlF_4_ and FLMyo6 in NF/high salt. **(C)** Rg of FLMyo6 WT, L926Q and SAHmimic determined by SEC-SAXS experiments in the presence of ADP.AlF_4_ and FLMyo6 in NF/high salt. Rg values were extracted from linear fits of the Guinier plots shown in Sup Fig. 1C using primusqt (ATSAS suite^39^). **(D)** Actin-activated ATPase rate of FLMyo6 WT, L926Q, SAHmimic and MD^Ins2^ (n=6).

### Structural model of the Myo6 off-state

The distinct EM density for the Jo-In fusion clearly defines the position of the N- and C-termini of FLMyo6 and demonstrates how it can lock the off-state. We used available Myo6 crystallographic structures to build a model inside the 3D reconstruction (Fig. 1F, see details in Methods). By defining the orientation of the Lever arm, the model revealed that the flexible joint allowing back-folding must be localized around aa 912-918 prior to the SAH. The ~10 nm long SAH ends up close to the Myo6 N-terminus and Converter, where the rest of the Tail can also participate in stabilizing interactions. Importantly, only the pre-powerstroke structure of the Motor domain (which traps ADP.P_i_), not the Rigor (NF) structure leads to good model-to-map agreement (Fig. 1F, Sup Fig. 2E). We challenged this model by measuring affinities between Myo6 CBD^1035-end^ and Myo6 Head fragments (Table 1, Sup Fig. 3). The CBD binds to the Motor domain with low affinity, and the strongest interaction (K_D_ ~150 nM) was measured for MD^Ins2/IQ/3HB^. Removal of the IQ-3HB region (MD^Ins2^) reduces the affinity by 2-fold. Last, the interaction between Myo6 CBD and MD^Ins2/IQ/3HB^ drops from *K*_*D*_ ~150 nM to ~750 nM upon nucleotide removal (Table 1, Sup Fig. 3). These data indicate an interaction of the CBD with both the Motor domain and the Lever arm and highlight the importance of the nucleotide state for optimal interaction.

**Figure 3.**
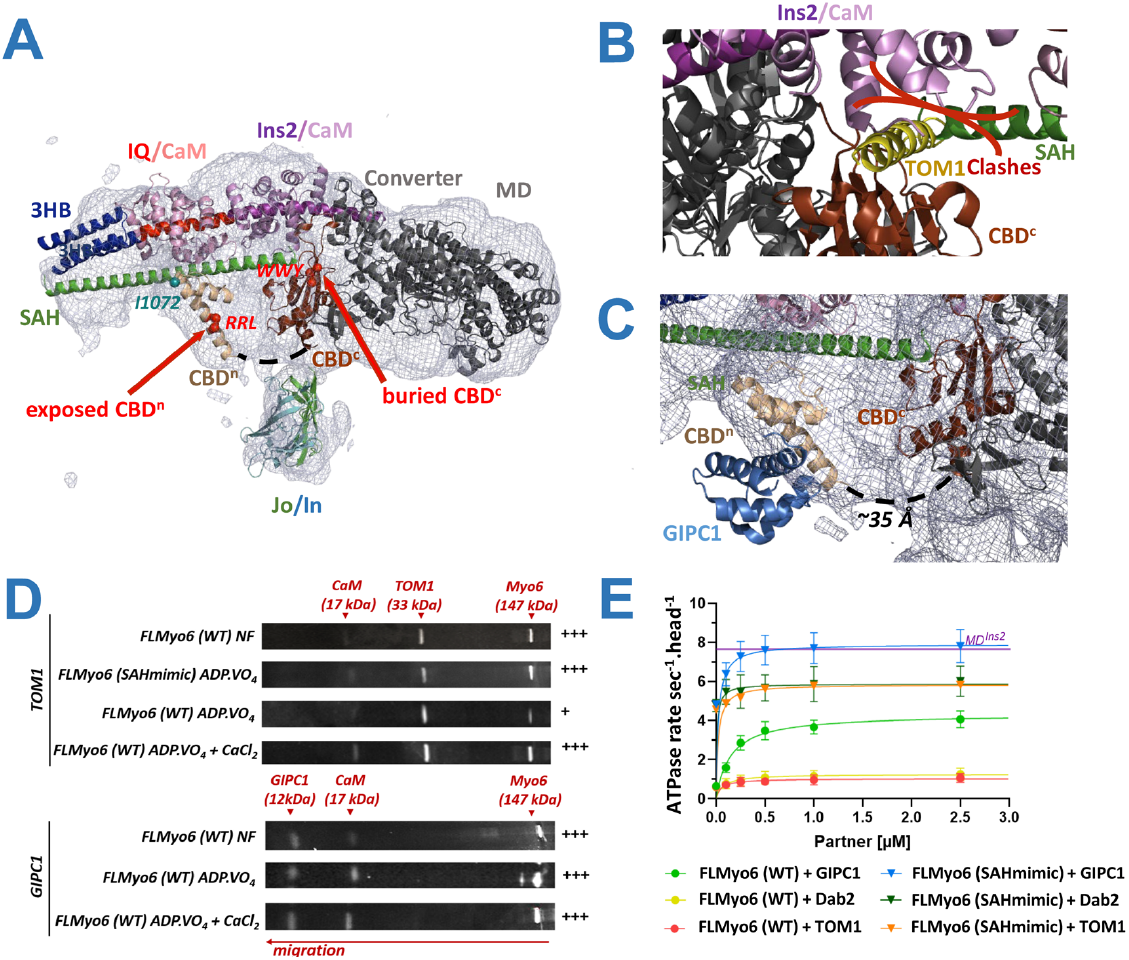
GIPC1 can bind to and activate the back-folded form of Myo6, while Dab2 and Tom1 canonly bind Myo6 once the motor has been primed open. **(A)** EM density for the Jo-Myo6-In (grey mesh) obtained by negative staining, as in Fig. 1H and Sup Movie 1. The WWY motif (red spheres) of CBD^c^ is buried. The CBD^n^ fragment (beige) (PDB: 5V6E,^34^) is positioned in the remaining, uninterpreted part of the density so that the RRL motif (red spheres) on CBD^n^ and the I1072 (blue sphere) that has been proposed to mediate interaction between ubiquitin and Myo6^41^ are both exposed. Note that no experimental model exists for 36 missing residues between the CBD^n^ and CBD^c^ (dashedlines). **(B)** Fitting of CBD^c^-TOM1 structure (PDB: 6J56^35^) with CBD^c^ (brown) in the model presented in Fig. 1H, TOM1 (yellow) needs to bind in a poorly exposed area and would clash with SAH (green) and CaM (lilac). **(C)** Fitting of CBD^n^ (beige)-GIPC1 (light blue) structure (PDB: 5V6E^34^) as for CBD^n^ alone. GIPC1 binding seems compatible with the back-folded conformation of Myo6 on a highly exposed binding site. **(D)** Anti-His pull-down assays. His-tagged Myo6 partners TOM1 and GIPC1 against FLMyo6 (WT) or (SAHmimic) revealed using SYPRO^42^ (Input and last wash pictured in Sup Fig. 5). Crosses: quantification of retained Myo6 (Image-Lab software, Bio-Rad) followed by stoichiometric normalization based on partner concentration (n=4). + means less than 10% Myo6 retained; +++ means more than 20% Myo6 retained. **(E)** ATPase rates of FLMyo6 (WT) and FLMyo6 (SAHmimic) with 40 µM F-actin and increasing concentrations of GIPC1, TOM1 or Dab2 (n=6). Purple line: ATPase rate of MD^Ins2^ at 40 µM actin (n=6) for reference.

To define the CBD region that interacts with the MD^Ins2/IQ/3HB^, we introduced four missense mutations (D1157V.Y1159D.D1161R.Q1163V: CBD^c^ loop mutant) in a conserved and exposed loop of the CBD (Fig. 1G). These mutations abolished the ability of the CBD to bind to MD^Ins2/IQ/3HB^, suggesting a key role of the CBD^c^, and this specific loop, in the interaction (Table 1). When this information is used to dock the CBD^c^, the Myo6 C-terminus is oriented towards the surface, close to the N-terminus consistent with our Jo-Myo6-In model (Fig. 1H, Methods).

### Auto-inhibition of Myo6 and hearing loss

The back-folded model predicts that a sharp kink occurs at the junction between the 3HB and the SAH (Fig. 1F). The N-terminus region of the SAH (aa 922-935) is thus positioned alongside the 3HB and could participate in the stabilization of the Myo6 off-state via apolar residues found in its atypical sequence (Fig. 2A). The importance of the sequence following the 3HB for back-folding was characterized using the previously published Myo6 (SAHmimic) mutant^17^, in which all apolar residues in the SAH were replaced by charged residues to match the i, i+4 alternance of a “perfect SAH sequence” (Fig. 2A). SEC-SAXS and SEC-MALS experiments indicated that FLMyo6 (SAHmimic) adopts an elongated conformation, even upon addition of an ADP.P_i_ analog, confirming the importance of the residues 922-935 for stabilization of the Myo6 off-state (Fig. 2B-C and Sup Fig. 4A).

**Figure 4.**
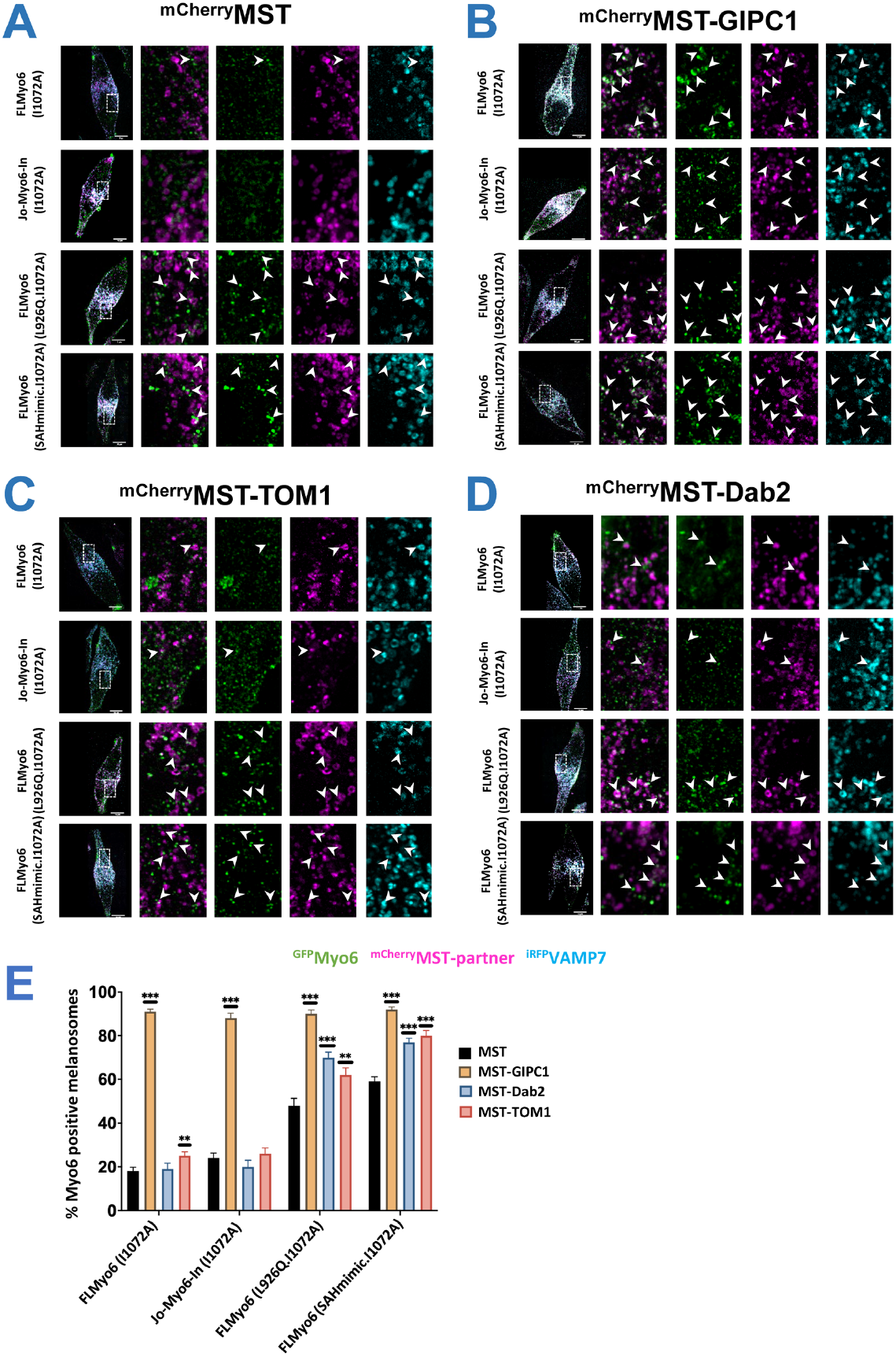
GIPC1 recruits Myo6 to melanosomes independently of Myo6 closure; Dab2 and TOM1 can only recruit Myo6 after the motor has been primed open. **(A)** Representative fixed MNT-1 cells co-expressing different ^GFP^Myo6 (I1072A), ^mCherry^MST and ^iRFP^VAMP7 constructs. **(B)** Representative fixed MNT-1 cells co-expressing different ^GFP^Myo6 (I1072A) constructs with ^mCherry^MST-GIPC1 and ^iRFP^VAMP7. **(C)** Representative fixed MNT-1 cells co-expressing different ^GFP^Myo6 (I1072A) constructs with ^mCherry^MST-TOM1 and ^iRFP^VAMP7. **(D)** Representative fixed MNT-1 cells co-expressing different ^GFP^Myo6 (I1072A) constructs with ^mCherry^MST-Dab2 and ^iRFP^VAMP7. **(A-D)** Green: Myo6 GFP; Cyan: ^irRFP^VAMP7; Magenta: ^mCherry^MST partner. From left to right: entire cell, 3 channels merged; 8x zoom on boxed region: ^GFP^Myo6 / ^mCherry^MST-partners merged, then individual channels. Scale bars: 10µm. Arrowheads: recruitment of Myo6 on melanosomes. **(E)** Myo6-positive melanosomes quantification of different ^GFP^Myo6 mutants when different ^mcherry^MST tagged partners are expressed (n=3, total cell number~30). Myo6-positive melanosomes are expressed in percentage and normalized to the total number of VAMP7-positive melanosomes. Cells were fixed 48h post-transfection then imaged and processed for quantification. Data are presented as the mean ± SEM. Significant stars: ***, P < 0.001; **, P < 0.01; *, P < 0.05, n.s., not significant (unpaired t test with Welch’s correction), for each ^GFP^Myo6 construct, significance of experiments with partners compared to the control without partner (in black on the graph).

Interestingly, a missense mutation present in this region of the SAH (L926Q) leads to deafness in humans^38^. Positioned away from the Motor domain or from Tail regions involved in recruitment (Fig. 2A), the effect of the mutation on Myo6 function had remained elusive. SEC-MALS (Sup Fig. 4A) and SEC-SAXS experiments (Fig. 2B-C) indicate that the L926Q mutation destabilizes the back-folded state. Both FLMyo6 (L926Q) and FLMyo6 (SAHmimic) mutants display higher ATPase rates (2.86 ± 0.12 and 4.83 ± 0.11 s^-1^.Head^-1^ respectively) than the wild-type (0.65 ± 0.08 s^-1^.Head^-1^) (Fig. 2D), which confirms the destabilization of the off-state.

We then investigated the impact of back-folding misregulation in the human pigmented melanoma cell line (MNT-1). Myo6 localizes to dot-like subdomains on the surface of pigmented melanosomes to promote membrane constriction and fission for the release of tubular carriers^21^. MNT-1 cells were transiently co-transfected with plasmids encoding (1) fluorescent components associated with pigmented melanosome ^iRFP^VAMP7^21,39^ and ^mCherry^MST^40^, and with (2) either FLMyo6 WT, SAHmimic, L926Q or Jo-Myo6-In, all fused to GFP. All ^GFP^Myo6 constructs localize as dots on melanosomes (Sup Fig. 4C), although at distinct levels (Sup Fig. 4B). The co-distribution of ^GFP^FLMyo6 (SAHmimic) or ^GFP^FLMyo6 (L926Q) with melanosomal components was greater than that of the ^GFP^FLMyo6 (WT) (~1.2 fold increase, p ≤ 0.001, Sup Fig. 4B). However, the co-distribution of ^GFP^Jo-Myo6-In with melanosomes was reduced ~3-fold compared to ^GFP^FLMyo6 (Sup Fig. 4B) and the associated cytosolic and diffuse fluorescent signal was more readily observed (Sup Fig. 4C).

Collectively, these data indicate that Myo6 auto-inhibition drastically reduces endogenous recruitment to melanosomes while impairment of Myo6 back-folding can result in over-recruitment. These results highlight the importance of the 3HB-SAH region for Myo6 auto-inhibition since the deafness L926Q mutation is sufficient for over-recruitment of the motor. Thus, destabilization of the off-state can lead to pathology.

### Differential binding and activation of FLMyo6 by distinct cellular partners

We next aimed at distinguishing whether partners can bind to FLMyo6 in the back-folded state and if binding depends on the specific binding site. Partners interacting either with the RRL motif (GIPC1^34^) or the WWY motif (TOM1^35^ and Dab2^14^) (Fig. 1A) were examined as our model suggests that in the FL off-state, the WWY motif of the CBD^c^ is buried and unavailable for binding (Fig. 3A-B). We first looked at the ability of ^His^GIPC1 and ^His^TOM1 to bind FLMyo6 using an anti-His pull-down assay on purified proteins, in conditions promoting either Myo6 back-folding (addition of ADP.VO_4_) or opening (NF, use of the SAHmimic mutant, or addition of Ca^2+^ as previously proposed^33,36^) (Fig. 3D, Sup Fig. 5). Both TOM1 and GIPC1 were able to retain Myo6 in conditions that favor Myo6 opening. In contrast, upon ADP.VO_4_ addition, the interaction of Myo6 with GIPC1 is maintained, but the interaction with TOM1 is weakened, suggesting that binding of TOM1 requires Myo6 opening.

**Figure 5.**
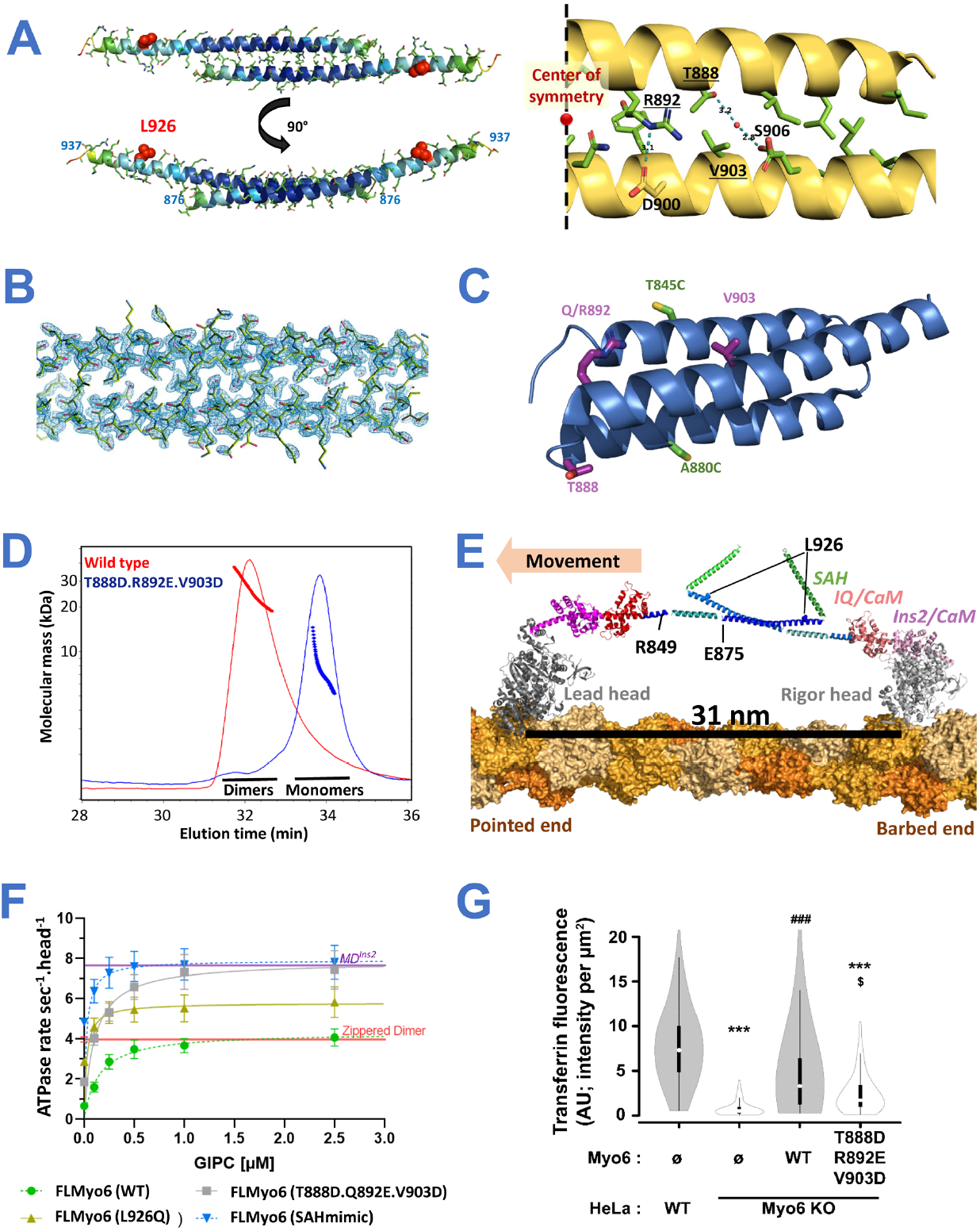
Myo6 can form an anti-parallel dimer through residues 875-940 which allow large steps. **(A) (Left)** X-ray crystallography structure of mouse Myo6 875-940 antiparallel dimer colored according to B-factor from 18.60 Å^2^ (dark blue) to 150.81 Å^2^ (red). **(Right)** Key residues for dimer stabilization. Apolar contacts are mediated by residues I878, L881, I885, T888, R892, I895, Q896, Y899, V903, E907, L909, L910, L913 (pictured in green). Dotted blue line: polar contacts. Residues mutated in our triple mutant (T888D.R892E.V903D) are underlined. **(B)** Close-up of the dimerization interface of Myo6 875-940 (crystallographic dimer) in the electronic density. **(C)** Triple helix bundle (PDB: 2LD3^43^) domain. T888, R892, V903, T845C and A880C pictured as sticks are surface residues. **(D)** SEC-MALS profiles of Myo6 875-940 WT (red) and T888D.R892E.V903D mutant (blue), following injection of 50 µl at 10 mg/mL in 10 mM Tris-HCl pH 7.5; 50 mM NaCl; 5 mM NaN_3_; 0.5 mM TCEP. WT elutes as dimers (32 µM at the peak) and T888D.R892E.V903D mutant elutes as monomers (43 µM at the peak). Measured molecular weight is heterogeneous due to the small size of the peptides through elution (low light scattering signal), but at the peaks it coincides with the expected masses. **(E)** Model of active FLMyo6 dimer (see Methods). **(F)** ATPase rates of FLMyo6 (WT) (green), FLMyo6 (T888D.Q892E.V903D) (grey), FLMyo6 (SAHmimic) (blue) and FLMyo6 (L926Q) (yellow) at 40 µM F-actin and increasing concentrations of GIPC1 (from 0 to 2.5 µM) (n=6). ATPase rates of MD^Ins2^ and zippered HMM dimer^12,44^ with no partner added (n=6) are plotted as purple and red thick lines (respectively) as references for monomeric and dimeric Myo6. **(G)** Fluorescence intensity of internalized transferrin was measured for each of the conditions after treatment with genistein (minimum 58 cells analyzed per condition). Expression of FLMyo6 (WT) construct restored transferrin uptake to wild-type levels, while the FLMyo6 (T888D.R892E.V903D), fails to rescue the defect. Data shown as violin plot (generated with BoxPlotR^45^) (p < 0.001, ANOVA; Tukey post-hoc comparisons: *** p < 0.001 vs WT, $ p < 0.05 vs Myo6 KO + FLMyo6 (WT)).

Next, we assessed the ability of GIPC1, Dab2 and TOM1 to stimulate the ATPase activity of FLMyo6 (Fig. 3E), and found that GIPC1 increases the Myo6 ATPase rate in a concentration-dependent manner, while addition of Dab2 or TOM1 has little impact. Note that partner affinities for Myo6 ^YFP^CBD are all in the submicromolar range (*i*.*e*., sufficient to ensure binding in our ATPase assays) (Sup Fig. 6). Lack of activation by TOM1 and Dab2 must thus be due to inaccessibility of the WWY motif in the back-folded FLMyo6. Interestingly, TOM1 and Dab2 increase the ATPase rate of the FLMyo6 (SAHmimic) mutant, as does GIPC1, which indicates that all partners can bind to and stabilize the unfolded state (Fig. 3E).

**Figure 6.**
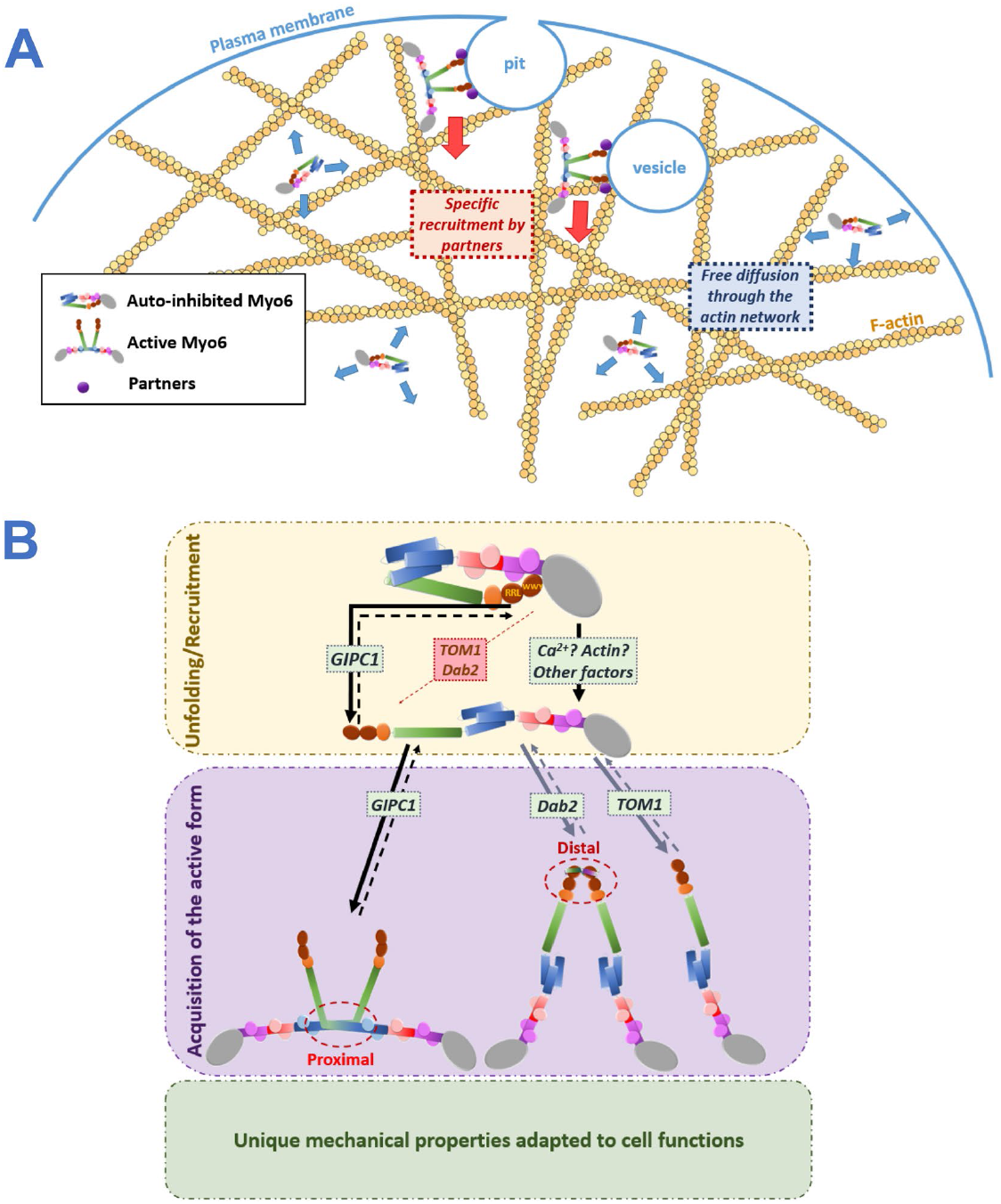
Importance of a folded monomer for regulation. **(A)** When auto-inhibited, Myo6 can diffuse across actin-rich regions without interacting with actin. Once recruited by a partner, Myo6 is activated and starts performing its cellular function. **(B)** Scheme representing possible activation mechanisms for Myo6. Myo6 MD is pictured in grey, Ins2/CaM in purple, IQ/CaM in red/pink, 3HB in blue, SAH in green, DT in orange and CBD in brown. Partner binding sites are labeled in garnet. The binding site (WWY) for Dab2 and TOM1 is blocked, preventing recruitment of Myo6 without a prior unfolding signal prior to unblock their binding. GIPC1 can bind the accessible RRL motif resulting in Myo6 recruitment and opening. Other signals can act as unfolding factors such as Ca^2+^, which can allow TOM1 to bind to Myo6. Such an activation cascade was previously proposed^36^. Once unfolded, Myo6 potentially acts as a monomer, as previously proposed^35^ upon TOM1 binding; or it can dimerize^29^ through proximal dimerization, as demonstrated in this study with GIPC1 binding; or it dimerizes through distal dimerization upon Dab2 binding^14^, which may lead to proximal dimerization.

In this context, we postulate that the RRL motif required for GIPC1 binding to CBD^n^ (Fig. 1A) must be exposed on the surface of the back-folded Myo6, as opposed to the WWY motif. The CBD^n^ fragment (*PDB: 5V6E*^34^) was thus positioned in the unexplained density lying in continuity to CBD^c^ in our Jo-Myo6-In 3D reconstruction (Fig. 3A-C, Sup Movie 1).

Collectively, these results demonstrate that not all partners can induce activation of Myo6. Partners Dab2 and TOM1 require another factor promoting Myo6 opening prior to their binding. In contrast, GIPC1 can directly activate FLMyo6, consistent with a previous study^33^.

### Assessing the specific recruitment of Myo6 to native organelles by distinct partners

If some partners can relieve Myo6 from auto-inhibition, we reasoned that artificial targeting of these partners to specific cellular membranes would lead to massive recruitment of Myo6. We thus decided to artificially drive GIPC1, TOM1 and Dab2 to melanosome membranes by fusing them to the melanosome-targeting tag (MST)^40^ (Sup Fig. 7) and verifying their ability to recruit either open (SAHmimic and L926Q) or locked (Jo-Myo6-In) Myo6. To do so with an optimized signal to noise measurement, we introduced the point mutation I1072A in our Myo6 constructs since it drastically reduces endogenous recruitment of Myo6 to the melanosomes (^GFP^FLMyo6 (I1072A), 3.7-fold reduction (p>0.0001) compared to ^GFP^FLMyo6; Fig. 4A; Sup Fig. 8A-B, 9A), while it is not part of the interface with either GIPC1 (Sup Fig. 6), TOM1 or Dab2. Hence, the I1072A mutation provides an easy way to reduce endogenous recruitment to melanosomes and offers a powerful tool to test the ability of distinct exogenous partners to recruit Myo6.

We transiently transfected MNT-1 cells with plasmids encoding for ^mCherry^MST-GIPC1, ^mCherry^MST-TOM1, or ^mCherry^MST-Dab2 for melanosome targeting. Co-transfection with plasmid encoding for ^GFP^Myo6 (I1072A), ^GFP^Jo-Myo6-In (I1072A), ^GFP^FLMyo6 (SAHmimic.I1072A) or ^GFP^FLMyo6 (L926Q.I1072A) provided a quantitative way to compare the ability of these partners to recruit Myo6 to specific organelles such as the melanosomes in cells (see Methods).

Expression of ^mCherry^MST-GIPC1 resulted in ~90% of Myo6-positive melanosomes for all the Myo6 constructs tested (Fig. 4B, E; Sup Fig. 9B), indicating that exogenous GIPC1 can recruit Myo6 to melanosomes independently of Myo6 being open or closed. In contrast, expression of ^mCherry^MST-TOM1 or ^mCherry^MST-Dab2 did not significantly increase the amount of ^GFP^Jo-Myo6-In (I1072A) positive melanosomes (p=0.5005 and p=0.344 respectively). This confirms the ineffectiveness of WWY partners in recruiting back-folded FLMyo6. Yet, their expression results in a 1.3/1.4-fold increase of melanosomes containing active Myo6 mutants ^GFP^FLMyo6 (SAHmimic.I1072A) and ^GFP^FLMyo6 (L926Q.I1072A) (p=0.003 or lower) (Fig. 4C-E; Sup Fig. 9C-D).

Interestingly, I1072A moderately affects the recruitment of Myo6 mutants impaired in auto-inhibition. Compared to ^GFP^FLMyo6 (SAHmimic) and ^GFP^FLMyo6 (L926Q), we observe reductions of 1.4 and 1.7-fold in the co-distribution with melanosome components for ^GFP^FLMyo6 (SAHmimic.I1072A) and ^GFP^FLMyo6 (L926Q.I1072A), respectively (Fig. 4A, 4E; Sup Fig. 4B-C), which are interestingly similar to the 1.6-fold reduction in recruitment observed for the CBD alone carrying the mutation I1072A (Sup Fig. 8C-D). We thus conclude that the I1072A mutation must reduce the affinity of the CBD for partner(s) responsible for Myo6 endogenous recruitment to melanosomes. In addition, the drastic reduction in ^GFP^FLMyo6 (I1072A) recruitment evidences the role of endogenous partners to promote Myo6 unfolding and indicates the major role of the I1072 residue in this process.

These results illustrate the importance of the recognition of the inactive state, and the distinct ways signaling factors can trigger association or activation of the back-folded state in a compartment for spatial and timely control of motor activity.

### A hinge that dimerizes

While we have demonstrated the key role of the sharp kink (hinge) at the 3HB/SAH junction for Myo6 auto-inhibition (Fig. 2; Sup Fig. 4), previous evidence by single molecule motility assays^12,17^ already suggested that this region is key for Myo6 proximal dimerization. Since Myo6 proximal dimerization might be critical for a number of functional properties, we wanted to elucidate the structure of the dimerization region.

SEC-MALS with six fragments derived from the 3HB/SAH junction (Sup Fig. 10A, C-E) indicated that a rather conserved region (aa 875-940) can self-dimerize with *K*_*DApp*_ of ~19 µM obtained from titration (Sup Fig. 10A, C). This minimal region corresponds to the last half of the 3HB (*i*.*e*., the 2^nd^ and 3^rd^ helix) and the first part of the SAH (Sup Fig. 10A-B). In contrast, no dimerization was observed when peptides included the whole 3HB domain, even when peak concentration of 30 μM was reached for the 834-955 peptide (Sup Fig. 10A, D). This data is consistent with previous findings indicating that proximal dimerization requires unfolding of the 3HB (^17^, Sup Fig. 11A).

Crystals of the 875-940 peptide diffracted to 2.1 Å resolution (Fig. 5A; Sup Table 1). Clear electron density for all residues from 876 to 937 indicates that they form an extended helix that dimerizes in an anti-parallel manner (Fig. 5B; Sup Fig. 12). This anti-parallel dimerization is stabilized by multiple apolar contacts involving 13 residues from each helix, and six polar contacts involving R892 with D900, and T888 with S906 (*via* a water molecule) (Fig. 5A; Sup Fig. 10B).

At the center of the interface, the structure highlights how residues T888, R892 and V903 contribute to the dimerization (Fig. 5A; Sup Fig. 10B). Three mutations (T888D.R892E.V903D) were introduced into the 875-940 peptide to assess their ability to disrupt proximal dimerization. Importantly, these residues were chosen on the surface of the 3HB so that the mutations would not disturb the bundle (Fig. 5C). (Note that residue 892 can be a Gln or an Arg depending on the species (Sup Fig. 10B) but both are compatible with the formation of the dimer). SEC-MALS confirms that the T888D.R892E.V903D mutant stays monomeric even up to 43 µM (peak concentration in the SEC-MALS experiment) while the WT counterpart is dimeric in similar conditions (Fig. 5D).

Finally, a model of the active dimeric configuration of Myo6 bound to F-actin including this crystal structure was built (Fig. 5E, see Methods). The inter-head distance is indeed compatible with the large (~30 nm) stepping previously reported when Myo6 walks processively^12,44^. Taken together, our results strongly support that proximal dimerization requires the formation of an extended anti-parallel coiled-coil, which can form following destabilization of the 3HB.

### GIPC1 promotes unfolding of the Myo6 monomer and proximal dimerization

We further characterized this proximal dimerization region and investigated the ability of partners to promote proximal dimerization of Myo6 using an actin-based ATPase assay. Such dimerization indeed leads to “gating”, *i*.*e*. coordination between the two Heads of the dimer that translates into slowing of ATP binding to the lead Head while the rear Head is attached^46^. This results in a 50% drop of ATPase rate per Head when Myo6 is dimerized compared to a monomer. The ATPase rate of zippered HMM (Myo6 truncated at R991 followed by a leucine zipper^47^ to create a constitutive dimer) in which gating has been characterized^46^ is indeed ~50% that of the monomeric MD^Ins2^ ATPase rate (Fig. 5F).

Indeed, upon addition of GIPC1, we found a ~50% reduction in the maximal ATPase activity per Head for FLMyo6 (WT) compared to MD^Ins2^ (Fig. 5F), consistent with GIPC1 promoting proximal dimerization of FLMyo6. In contrast, addition of GIPC1 to the FLMyo6 (SAHmimic) mutant is similar to that measured with MD^Ins2^, consistent with a role of GIPC1 in fully freeing the Motor domain from Tail inhibition upon stabilizing an extended, monomeric conformation. Importantly, the FLMyo6 (T888D.Q892E.V903D) exhibits ~2-fold higher maximal ATPase rate upon GIPC1 addition, consistent with loss of gating (Fig. 5F).

This additional evidence strongly validates the role of these residues in antiparallel proximal dimerization, and the role of this region in controlling motor mechanical properties. Furthermore, we demonstrate for the first time that proximal dimerization (involving 3HB unfolding) can be triggered upon GIPC1 binding.

### The L926Q deafness mutation indirectly impairs proximal dimerization

Interestingly, when we used GIPC1 to activate the FLMyo6 (L926Q) construct (Fig. 5F), the maximal ATPase activity that we found was intermediate between monomeric and dimeric FLMyo6. Since our proximal dimerization structure indicates that the L926Q missense mutation does not impact the anti-parallel coiled-coil region itself (Sup Fig. 11B), and since we found that 3HB unfolding is essential for proximal dimerization, we hypothesized that L926Q impairs Myo6 dimerization by perturbing the unfolding of the 3HB. This was previously reported for the FLMyo6 (SAHmimic) mutant^17^.

To monitor 3HB unfolding, we introduced cysteines at two positions of the 3HB surface (T845 and A880), for tetramethylrhodamine (TMR) labelling (Fig. 5C, Sup Fig. 11A). As previously described^18^, a low TMR fluorescence ratio is found when 3HB is folded (fluorescence quenching due to stacking of the rhodamine rings; MD^Ins2/IQ/3HB^ T845C, A880C). This value increases upon 3HB opening (Sup Fig. 11A), and a high TMR fluorescence ratio indicative of 3HB unfolding has been reported for the Myo6 zippered HMM T845C, A880C^18^. Introducing the L926Q mutation in this zippered construct led to an intermediate fluorescence intensity, indicating limited unfolding of the 3HB for the deafness mutant compared to control (Table 2). This suggests a role for the L926Q mutation in limiting the conformational changes of the 3HB required for dimerization, in addition to its effect in destabilizing the off-state.

**Table 2.** L926Q stabilizes the 3HB. Fluorescence observed by TMR labeling of one or two cysteine residues inserted into the three-helix bundle of monomers (MD^Ins2/IQ/3HB^) and dimers (zippered HMM). Fluorescence was analyzed by a ratio of the emission values to that of the absorption values for each construct from four independent measurements (n=4). Mean values (±SD) are reported. The molar ratio was calculated by comparing the myosin concentration to the concentration of the incorporated TMR.

| Construct | Fluorescence Ratio without Actin + ATP | Fluorescence Ratio with Actin + ATP | Molar Ratio of Labeling per Myosin Head |
| --- | --- | --- | --- |
| MD<sup>Ins2/IQ/3HB</sup> T845C | 256.1 ± 24.4 | 243.8 ± 14.5 | 1.03 |
| MD<sup>Ins2/IQ/3HB</sup> T845C, A880C | 22.5 ± 5.8 | 22.6 ± 7.2 | 2.10 |
| zippered HMM A880C | 238.2 ± 23.7 | 232.3 ± 30.4 | 1.11 |
| zippered HMM T845C, A880C | 206.5 ± 19.6 | 214.3 ± 12.6 | 2.22 |
| Zippered HMM T845C, A880C, L926Q | 147.53 ± 30.4 | 164.6 ± 18.7 | 2.14 |

### Importance of proximal dimerization in cells

To further demonstrate that proximal dimerization of Myo6 occurs via the anti-parallel coiled-coil seen in our structure, we compared the ability of FLMyo6 (WT) and FLMyo6 (T888D.R892E.V903D) to rescue Myo6-mediated transferrin uptake^17^ in HeLa cells whose Myo6 was knocked out for Myo6 by using CRISPR/Cas9 (Sup Fig. 13A). FLMyo6 (WT) and the FLMyo6 (T888D.R892E.V903D) were transiently expressed, and the transferrin internalized during a 10 min pulse was quantified. As summarized in Fig. 5G and Sup Fig. 13B, expression of the T888D.R892E.V903D mutant, unable to form the proximal dimer, profoundly decreases the rate of uptake of endocytic vesicles, providing evidence for the need of proximal dimerization to optimize Myo6 function during endocytosis. Furthermore, this also strengthens the evidence that proximal dimerization occurs via an anti-parallel coiled-coil as depicted in our crystal structure.

## Discussion

Despite the significance of controlling where and when myosin motors generate forces and move cargoes in cells, careful investigation of how the function of myosin motors is regulated has only been performed for a few classes of myosin^2,3,48,49^, and most extensively for Myo2. The results of this study highlight the importance of regulated inhibition of the Myo6 motor until it reaches its target in a cell and it is activated. Myo6 must cross actin-rich regions in order to diffuse and reach its binding partners which selectively activate motor activity (Fig. 6A). If the motor was not blocked from interacting with and cycling on actin, Myo6 would bind to actin filaments throughout the cell, retarding diffusion to its target sites at the cell membrane. The fact that the L926Q mutant disrupts the folding and regulation of Myo6 (Fig. 2) and causes deafness in humans^38^ attests to the critical need for the regulation of this class of myosin motors.

Our structural and functional studies provide a more precise model to account for the interactions stabilizing Myo6 back-folding (Fig. 1H, Table 1). Among the major differences compared to previous models^7,36^, we show that **(1)** ADP.P_i_ bound to the Motor domain is essential to lock Myo6 in its back-folded state (Fig. 1B, Table 1); **(2)** back-folding involves a specific loop of the CBD^c^ (Table 1), which was previously predicted to be external to the folded complex by Alphafold^50,51^ (Sup Fig. 14); and **(3)** the 3HB/SAH junction acts as a critical hinge to control the equilibrium between on/off states of the motor (Fig. 1G-H, Fig. 2). Earlier studies of the folded monomers^25,28,29,36^ focused only on interactions within a full-length construct in the absence of nucleotide or within a Motor-less construct, and thus did not fully represent what is happening with the full-length Myo6 monomer saturated with nucleotide.

Intriguingly, a single amino acid change (L926Q) causes deafness^38^ and is in fact sufficient to destabilize the back-folded monomer (Fig. 2). This SAH mutation flanks the hinge region that we identified as essential for the off-state of this motor. To further investigate the impact of Myo6 back-folding in myosin recruitment, we used the FLMyo6 (L926Q) and FLMyo6 (SAHmimic) mutants to probe their impact on Myo6 recruitment on melanosomes. What was observed (Sup Fig. 4B-C) was that both constructs lead to greater recruitment than the FLMyo6 (WT). This is not a gain of function, but rather a loss of regulation as, **(1)** the normal cellular control over the spatial and temporal recruitment of Myo6 has been lost, and **(2)** fluorescence quenching assays show that both SAHmimic^17^ and L926Q (Table 2) mutations impair proximal dimerization and thus Myo6 function. Deafness due to the L926Q mutation in humans may therefore result from the inability of Myo6 monomers to reach their target sites in hair cells due to loss of folded regulation.

We next examined the ability of some of the Myo6 binding partners that recognize different regions of the CBD to induce unfolding and recruitment of Myo6. Folding not only prevents cycling of the motor on actin until the cellular target has been reached, but as shown by our actin-activated ATPase (Fig. 3D), pull-down (Fig. 3E) and recruitment assays on melanosomes (Fig. 4), the folding can also prevent interaction with a subset of cellular partners until unfolding occurs, by either a different class of partners, or potentially by a spike in cellular Ca^2+^ concentration^33,36^ or PIP_2_ recognition^43,52^. We thus propose a model of the folded off-state of FLMyo6 in which the GIPC1 binding site is available for binding, while the TOM1/Dab2 site is masked (Fig. 3A-C). Interestingly, this demonstrates that not all partners are equivalent in their potential for binding the auto-inhibited form of the motor and to activate Myo6.

While TOM1 and Dab2 cannot trigger Myo6 initial unfolding, once bound they prevent the formation of the off-state due to their incompatibility with it, as previously proposed^25,28^. Depending on the nature of the partner and its distribution, binding will activate the Myo6 motor and could drive either proximal dimerization (Fig. 5F), distal dimerization^14,15^ or maintain an activated, monomeric form^35^. Taken together, these results suggest unique roles for partners not only in Myo6 localization, but also in the control of Myo6 activation and function (Fig. 6B).

Once unfolded, binding to its cargo brings two unfolded Myo6 monomers into close apposition, favoring its dimerization ^12,14,15,17,18,25^. The experiments summarized in Fig. 5 provide the previously unknown structure of the proximal dimerization region. We present both *in vitro* and cellular evidence in support of the structure. This structure reveals the dimerization region to be an anti-parallel coiled-coil, as for Myo10^9,11,13^. Mutations of three of the amino acids that stabilize this coiled-coil structure (T888D.Q892E.V903D) abolish dimer formation in *in vitro* assays, but with no impact on back-folding of the monomer (Fig. 5F). Furthermore, introduction of this Myo6 triple mutant into cells fails to rescue endocytosis (Fig. 5G), providing evidence for the need of proximal dimerization to optimize this cellular function of Myo6.

Myo7A, Myo10 and Myo6 exist as folded monomers in cells until they are activated and recruited by their partners. Formation of an antiparallel dimer may be the mode of dimerization for the three classes that appear to undergo this folded monomer to dimer transition. The structure of the active form of Myo6 has been a long-debated issue ^13,15,17,11^, which is resolved by our structure for the proximal dimerization region. As shown in Fig. 5E, Myo6 is unique in that its antiparallel coiled-coil and Lever arm in the dimer are derived from unfolding of a 3HB, with contribution of the SAH. The resulting Lever arm formed by the CaM binding region and the unfolded 3HB (half of which contributes to the coiled-coil) is sufficiently long to account for the ability of Myo6 to take steps that average ~30nm on actin^12,44^. These findings provide a structural framework that can be applied to understanding how motors are recruited and how partners influence motor functions in cells.

## Methods

### Constructs cloning, expression and purification

Details on protein cloning and purification can be found in Supplementary Methods.

### SEC-SAXS

SAXS data were collected on the SWING beamline at synchrotron SOLEIL (France)^53^ in HPLC mode at λ = 1.0332150494700432 Å using a Dectris EIGER-4M detector at a 2 m distance. Protein samples were injected at 0.1 mL/min on Superdex 3.2/300 column pre-equilibrated in 20 mM HEPES; 200 mM NaCl; 2 mM MgCl_2_; 1 mM NaADP; 1 mM AlF_4_, 0.1 mM EGTA, 1 mM DTT; or 20 mM HEPES; 400 mM NaCl; 2 mM MgCl_2_, 0.1 mM EGTA, 1 mM DTT; pH 7.5 prior to data acquisition in the SAXS capillary cell. 150 frames of buffer scattering (before the void volume), then 719 frames of elution sample scattering were collected. Exposure time was 1990 ms/frame. Images were processed using the Foxtrot 3.5.10-3979^53^ developed at the SOLEIL synchrotron: buffer averaging, buffer substraction from the corresponding frames at the elution peak, and sample averaging were performed automatically. Further data analysis to obtain Rg, I(0), Dmax and molecular weight estimation was done with PRIMUS from ATSAS suite^54^. Dimensionless Kratky plot ((q.Rg)^2^.I(q)/I(0) versus q.Rg) was generated using Microsoft Excel based on I(0) and Rg values found with PRIMUS. 20 envelopes were generated independently with GASBOR^55^ and averaged with DAMAVER^56^.

### SEC-MALS

For SEC-MALS analysis, samples were injected in a Superdex 200 10/300 Increase (GE Life Sciences) previously equilibrated in the corresponding buffer, and developed at 0.5 ml/min. Data collection was performed every 0.5 sec with a Treos static light scattering detector, and a t-Rex refractometer (both from Wyatt Technologies). Concentration and molecular mass of each data point were calculated with the software Astra 6.1.7 (Wyatt Technologies).

### Microscale Thermophoresis measurements between Myo6 Tail and Head

Microscale thermophoresis experiments were performed on a Monolith NT.115 system (NanoTemper Technologies) using YFP-fusion proteins.

The non-fluorescent protein was first treated with 0.5 mM EGTA (+/-2 mM MgADP; 2 mM Na_3_VO_4_ for some experiments); then dialysed against 20 mM Hepes pH 7.5, 50 mM NaCl, 2.5 mM MgCl_2_, 1 mM TCEP and 0.05% (v/v) Tween-20 (+/-2 mM MgADP; 2 mM Na_3_VO_4_ for some experiments). Two-fold dilution series (16 in total) of the non-fluorescent protein (« Head » sample) were performed at 25° in the same buffer. The YFP-fused partner was kept at a constant concentration of 100 nM. The samples were loaded into premium capillaries (Nanotemper Technologies) and heated for 30 sec at 60% laser power. All experimental points were measured twice. The affinity was quantified by analyzing the change in thermophoresis as a function of the concentration of the titrated protein using the NTAnalysis software provided by the manufacturer.

### Microscale Thermophoresis measurements with Myo6 partners

Two-fold dilution series (16 in total) of the non-fluorescent protein (Myo6 partner) were performed at 25°C in the MST buffer: 20 mM Bis-Tris pH 6.5, 100 mM KCl, 1 mM DTT and 0.05% (v/v) Tween 20. The YFP-fused partner was kept at a constant concentration of 100 nM. Microscale thermophoresis experiments were then performed in similar conditions as above in 20 mM Bis-Tris pH 6.5, 100 mM KCl, 1 mM DTT and 0.05% (v/v) Tween 20. Capillaries were heated for 30 sec at 50% laser power.

### ATPase assays

ATPase assays were performed as previously described^57^. ATPase rate determined from 2-3 preps with 2-3 independent assays per prep. F-Actin was used at 40 µM and Myo6 at 150 nM with 2.5 µM additional CaM in all our experiments. The experiments were all carried out in 10 mM MOPS pH 7.0; 10 mM KCl; 1 mM DTT; 1 mM MgCl_2_; 1 mM EGTA.

### P_i_ release experiments

Actin-activated phosphate release from Myo6 constructs was monitored using the 7-diethylamino-3-((((2-maleimidyl)ethyl)amino)carbonyl) coumarin labeled phosphate binding protein (MDCC-PBP) as in ^57^. Experiments were conducted under single turnover conditions where the substrate was rate-limiting. Actin-stimulated phosphate release was measured in a double-mixing experiment where Myo6 was mixed with a substoichiometric concentration of ATP, aged for 30 seconds, and subsequently mixed with actin and MDCC-PBP (final reaction conditions: 0.5 μM Myo6, 0.25 μM ATP, 14 μM actin, 5 μM MDCC-PBP). Fluorescence was monitored by excitation at 425 nm, and emitted light was collected through a 455 nm long pass glass filter. Myosin solutions were clarified by centrifugation (550k x gmax, 10 minutes, 4°C) after thawing and buffer exchange.

### Anti-His Pull-down assay

FLMyo6 SI (WT) or (SAHmimic) were used alone or mixed with partner GIPC1 (*His-rTEV-GIPC1 255-end*) or TOM1 (His-TOM1 207-492) in a ratio (1/1) (10µM) and 1 µM of extra Calmodulin was added in a total volume of 20 µL. The input was incubated with 40 µL of Ni^2+^ beads from cOmplete column (Roche), which were previously equilibrated either in ADP.VO4 Buffer (10 mM HEPES pH 7.5; 100 mM NaCl; 5 mM NaN_3_; 1 mM MgCl_2_; 0.1 mM TCEP; 1 mM NaADP; 1 mM Na_3_VO_4_; 0.1 mM EGTA; 4 mM imidazole) or ADP.VO4-CaCl_2_ Buffer (10 mM HEPES pH 7.5; 100 mM NaCl; 5 mM NaN_3_, 1 mM MgCl_2_, 0.1 mM TCEP, 1 mM NaADP; 1 mM Na_3_VO_4_; 4 mM imidazole; 1 mM CaCl_2_) or NF Buffer [10mM HEPES pH 7.5; 100 mM NaCl; 5 mM NaN_3_; 1 mM MgCl_2_; 0.1 mM TCEP; 0.1 mM EGTA; 4 mM imidazole]. All steps were performed at 4°C. Beads were washed by centrifugation after 1 hour of gentle agitation. Bound proteins were eluted in 600 mM imidazole in the corresponding buffer.

### Electron Microscopy

Purified Jo-Myo6-In at 50 µg/ml in 10 mM HEPES; 80 mM NaCl; 1 mM MgCl_2_; 0.1 mM TCEP; 0.1 mM ADP; 0.2 mM Na_3_VO_4_; 0.1 mM EGTA, pH 7.5 was transferred to Carbon Film 300 mesh copper grids (Electron Microscopy Sciences), then stained with 2% uranyl acetate. A total of 284 images were collected with a 200 kV Tecnai G2 microscope under low dose condition with a 4Kx4K F416 TVIPS camera at 0.213 nm/px and treated with the software CryoSPARC^58^. Following CTF determination, template picking was carried out using an initial set of 100 manually picked particles. The resulting 711,671 particles were submitted to a few rounds of 2D classification from which 93,293 particles were selected. These were used in the ab-initio reconstruction that produced the map at 17 Å resolution (FSC).

### Model of the Myo6 off-state

We first positioned the Motor domain-Lever arm (residues 1-917) in the Jo-Myo6-In maps from SAXS and EM. Best model-to-map agreement was obtained with PDB 4ANJ^59^ (Motor domain and insert-2/Ca^2+^-CaM with ADP.P_i_ analog bound, in PPS state). The Lever arm from PDB 3GN4^18^ was then superimposed to PDB 4ANJ by using the insert-2/Ca^2+^-CaM region, present in both structures, as reference. In the negative staining reconstruction, Jo-In PDB 5MKC^37^ was placed in the distinct density that corresponds to it, as expected, with the N- and C-termini pointing towards the center of the main density body occupied by Myo6. The structure of the C-terminal half of the CBD from PDB 3H8D^14^ was placed according to structural and biochemical restrictions as follows: (1) the CBD^c^ C-terminus must be near the N-terminus of the fusion protein In; (2) residues D1157, Y1159, D1161 and Q1163 are in contact with the MD^Ins2^; (3) there is still density to be filled close to the N-terminus of the CBD^c^, that can be filled by CBD^n^. (Note that this proposed position is opposite to that currently predicted by Alphafold^50,51^ for uniprot entry : Q9UM54, due to lack of data for the intermolecular interactions when that model was built). At last, the NMR structure of the SAH domain (residues 919-998; PDB: 6OBI^60^) was accommodated in the density. This density is narrow up to residue ~955 and then becomes much larger to account for the rest of the model, in which no distinct subdomain can be identified. Thus, our current model lacks the distal Tail (a compact domain of 3 nm in diameter^7^) and CBD^n^, for which there seems to remain enough density to be fitted. Model and figure were prepared with Pymol^61^.

### MNT-1 cell transfection

MNT-1 cells were cultured in DMEM supplemented with 20 % FBS, 10 % AIM-V medium, 1 % sodium pyruvate, 1 % nonessential amino acids, and 1 % penicillin-streptomycin. For plasmid transfection, 400 000 MNT-1 cells were transfected using nucleofection (NHEM kit, Lonza) on Amaxa device 2 (program T20) with 1.5 µg of ^iRFP^VAMP7 plasmid; 1 µg of ^mCherry^MST plasmid and 3 µg of ^GFP^FLMyo6 plasmid. After transfection cells were seeded in fluorodish containing 1 mL RPMI medium, then 1 mL of complete MNT-1 medium supplemented by 10 % FBS was added 6 h post-transfection. Medium was changed 1-day post-transfection by complete medium then cells were fixed with 4 % PFA at 48 h post-transfection. Cells were stored in the dark at 4 °C in PBS medium until imaging. Fluorescence intensity of each ^mCherry^MST construct was analyzed to ensure equivalent expression levels between the different partners (Sup Fig. 15).

### Super resolution imaging and analysis

Samples were imaged in fluorodish using a 100x/1.4 NA oil immersion objective on an inverted Spinning disk confocal microscope (Inverted Eclipse Ti-E Nikon, Spinning disk CSU-X1, Yokogawa) equipped with a Photometrics sCMOS Prime 95B Camera (1200 × 1200 pixels). Z images series were acquired every 0.2 µm. Images were processed with a Live super Resolution module (Live-SR; Gataca systems) based on structured illumination with optical reassignment technique and online processing leading to a two-time resolution improvement^62^. For the figures, Z maximum projection and a substract background (50 pixels) were applied on SR images using the FIJI software. Analysis was done on raw SR images. Melanin pigments (black spots) were automatically detected in a defined region of interest (ROI) (here, cell outlines that were manually drawn) in BrightField images by creating a MIN-intensity z-projection and considering the lowest values, defined using the ‘Find Maxima’ function of Image J/Fiji and whose spatial coordinates were recorded. To quantify the percentage of melanosomes containing ^iRFP^VAMP7 / ^mCherry^MST / ^GFP^Myo6 proteins at the membrane, additional ROIs centered around each individual detected pigment were generated whose size was defined (0.350 µm diameter). Then, for each detected brightfield spot, an additional automatic detection in the fluorescent channel(s) of interest was performed by creating a MAX intensity z-projection in the ROI around the pigments and considering the highest values. Detected pigments were considered positive for the marker of interest above a threshold (defined by Triangle’s automatic thresholding method, calculated on the MAX intensity projection, or manual thresholding in the case of cells expressing the lowest GFP-Myo6), and the percentage of which was calculated. Pigments that were automatically detected very close to each other (within 4 pixels in XY and 2 pixels in Z) and that had overlapping ROIs were automatically removed and eliminated from the analysis to avoid data duplication. Moreover, automatically detected pigmented that were negative for ^iRFP^VAMP7 and/or ^mCherry^MST fluorescent signal were excluded from the analysis because not considered as pigmented melanosomes (positive for membrane-associated components). For each cell, a percentage of Myo6 positive melanosome was calculated and normalized to the total number pigmented melanosomes (co-positive for pigment and ^iRFP^VAMP7 and/or ^mCherry^MST).

### Proximal dimer Crystallization, Data Collection, and Structure Determination

The Myo6 875-940 construct was crystallized by hanging drop vapor diffusion at 290 K by mixing 1 µL of 9.8 mg/mL protein solution with 1 µL of reservoir solution (27% PEG 4000, 10 mM MgCl_2_ and 0.2 M imidazole / malate, pH 6.0). Crystals grew spontaneously as rods 1 to 7 days after. After additional 3 weeks, they were cryo-cooled in liquid nitrogen in a solution containing 28% PEG 4000, 10 mM MgCl_2_, 0.2 M imidazole / malate, pH 6.0, and 27% ethylene glycol. One exploitable X-ray dataset was collected at the Proxima 1 beamline (Synchrotron Soleil, Gif-Sur-Yvette) and processed with Autoproc^63^. Diffraction limits after treatment with Staraniso^64^ with cut-off of 1.2 I/sI were 2.566 Å in two directions, and 2.077 Å^64^ in one direction. Initial structure factors were obtained by molecular replacement with Phaser^65^ using a helix comprised of 30 serine residues as search model. Initial sequence attribution was obtained with Phenix AutoBuild^66^, followed by several cycles of iterative edition with Coot^67^ and refinement with Buster^68^. Resolution was automatically cut by Buster to 2.2 Å based on model-map cross-correlation. The dimer is defined by one of the 2-fold symmetry axis, with crystal contacts between the N-terminus of one dimer and C-terminus of neighboring dimers (Sup Fig. 12B-C, Sup Movie 2). When the carbons are colored according to B-factors (Fig. 5A), the lowest values are found between residues 885 and 913, suggesting that the dimerization interface is comprised within those boundaries.

### Model of the Myo6 proximal dimer

The cryoEM structure of Myo6 bound to actin (PDB: 6BNP^69^) was used as basis for placing two Myo6 Motor domains in rigor and PPS states (grey) at the desired 31 nm distance (Fig. 5E). On each side, the N-terminus of the Lever arm bound to two light chains (pink, PDB: 3GN4^18^) was aligned to the corresponding residues in the Converter. The crystallized dimerization domain (blue) was then placed with minimal distance from the two Lever arms, leaving a gap of 4.6 nm from each side. This gap needs to be filled by a stretch of 26 amino acid residues that would make 3.9 nm if in a theoretical helix, or up to 9.9 nm if fully extended. SAH (green) (PDB: 6OBI^60^) was connected to the C-terminus of the dimerization region *via* a putative kink. Model and figure were prepared with Pymol^61^.

### Bundle Unfolding Assay

As previously described^18^, cysteine residues were introduced to replace T845 and A880 in Myo6-917 and Myo6-991-GCN4 constructs with no reactive cysteines. Control constructs contained one reactive cysteine, T845C. One mg of each protein was labeled with a 10-fold molar excess of TMR 5-iodoacetamide (5-TMRIA; Anaspec, San Jose, CA) per cysteine (from a stock concentration of 20 mM in dimethylformamide) at 4°C for 1–3 hr. Unbound rhodamine was removed by gel filtration and overnight dialysis. Absorption spectra were measured in a HP Diode Array Spectrophotometer, and fluorescence spectra were obtained in a PTI QM3 luminescence spectrofluorometer. The excitation and emission spectra were measured at 552 and 575 nm, respectively.

### Generation of Myo6 null HeLa cells

The Myo6 gene of HeLa cells (ATCC CCL-2) was inactivated by CRISPR/Cas9 gene editing approach. Briefly, HeLa cells were transfected using a combination of three human Myo6 CRISPR plasmid variants (Santa Cruz Biotech SC-401815) – each driving expression of Cas9, GFP and one of the following human Myo6-specific 20-nucleotide gRNAs (5’-3’: taatatcaaagttcgatata, acattctgattgcagtgaatc, ccaagtgtttcctgcagaag). Clones of transfected HeLa cells were selected on the basis of GFP fluorescence, and PCR of isolated DNA using primers flanking the targeted genomic sequences. Loss of Myo6 expression was confirmed by western blot.

### Transferrin Endocytosis Assay

Normal and Myo6 null HeLa cells were grown in multi-well tissue culture plates on coverslips coated with rat collagen I (Corning). FLMyo6 (WT) or FLMyo6 (T888D.Q892E.V903D) were tagged with a C-terminal mApple for identification of expressing cells. Transfections were performed using the X-tremeGENE 9 DNA transfection reagent (Sigma-Aldrich) following the manufacturer’s instructions. Cells were serum starved, but otherwise maintained in normal growth conditions – at 37 °C with 5% CO_2_, by incubation in serum-free Dulbecco’s modified Eagle’s medium for 2.5 h. Serum-free medium was supplemented with genistein (600 µM, Cayman Chemical Company) to inhibit caveolae-mediated uptake of transferrin following Myo6 depletion as previously reported^17^. During the final 10 min of serum starvation, Alexa fluor 488-conjugated transferrin (ThermoFisher) was added to the culture medium at 25 µg/ml. Following serum starvation, plates were placed on ice and washed twice with 10 mM HCl and 150 mM NaCl to remove cell surface-bound transferrin. The cells were fixed with ice-cold 4 % paraformaldehyde for 20 min and stained with rabbit anti-dsRed antibody (Takara Bio) and Alexa fluor 568-conjugated anti-rabbit secondary antibody to identify cells expressing mApple-tagged Myo6. Image acquisition was performed with a Leica Application Suite X software on Leica TSC-8 confocal system using a 40X oil immersion objective lens (n.a. = 1.3). Transferrin uptake was determined using ImageJ software: the total transferrin-conjugated fluorescence intensity from sum slice projections of individual cells was subsequently normalized by cell size. Comparative samples were stained, imaged, and processed simultaneously under identical conditions. Data were subjected to one-way analysis of variance with Tukey post-hoc comparison of individual groups to determine statistical significance.

## Supporting information

Supplementary informations

## Data availability

The atomic model of the Myo6 proximal dimer is available on the PDB^70^ under the accession code 8ARD. The other data supporting the findings of this study are available from the corresponding author upon request.

## Acknowledgments

The authors greatly acknowledge the Cell and Tissue Imaging (PICT-IBiSA) – Institut Curie, member of the French National Research Infrastructure France-BioImaging (ANR10-INBS-04) and in particular Anne Sophie Macé for her help in the quantification of the results; the CurieCoreTech Mass Spectrometry Proteomics and Recombinant protein; the beam line scientists of PX1 and SWING (SOLEIL synchrotron) for excellent support during data collection; Pierre-Damien Coureux for preliminary experiments in negative staining electron microscopy; Margaret Titus for her help with Jo-Myo6-In cloning and comments on the manuscript; Virginie Ropars for her help in SAXS data optimization, Guillaume Jousset for its advises in biochemistry and Christopher Toseland for accepting to provide us with Dab2 DNA.

## Author contributions

A.H. and H.L.S. designed and directed the research. L.C., V.J.P.-H., K.J.P. and C.D. were involved in project management. H.S., X.L., L.H., L.C., T.L., C.K., L.V., A.D., E.F., P.H., V.J.P.-H., D.M. cloned, produced and purified constructs. L.C., T.L., C.K., H.S., L.H., A.D., L.V., P.H., V.J.P.-H., D.M. performed biophysical and biochemical assays. C.K. performed grid optimization for negative staining EM that were collected by ADC. D.L. provided access to the 200kV microscope. C.K. analyzed the negative staining data and performed the 3D reconstruction. H.S. characterized the limits of the proximal dimerization region, crystallized it and C.K. solved the structure. F.M., Y.I.L. and K.J.P. performed the cell-based assays. L.C., C.K., A.H., H.L.S., F.M., Y.I.L., H.S., L.H., A.D., L.V., K.J.P. and V.J.P.-H. analyzed the data. J. S.-C. provided technical help in quantification of the cell-based assays. L.C., A.H., C.D., C.K. and H.L.S. wrote the initial version of the paper, with the help of F.M. and V.J.P.-H. and all authors reviewed it. A.H. and H.L.S. provided fundings.

## References

1. Myosins: A Superfamily of Molecular Motors. vol. 1239 (Springer International Publishing, 2020).

2. Heissler, S. M., Arora, A. S., Billington, N., Sellers, J. R. & Chinthalapudi, K. Cryo-EM structure of the autoinhibited state of myosin-2. Sci Adv 7, eabk3273 (2021).

3. Niu, F. et al. Autoinhibition and activation mechanisms revealed by the triangular-shaped structure of myosin Va. Sci Adv 8, eadd4187 (2022).

4. Robert-Paganin, J., Auguin, D. & Houdusse, A. Hypertrophic cardiomyopathy disease results from disparate impairments of cardiac myosin function and auto-inhibition. Nat Commun 9, 4019 (2018).

5. Umeki, N. et al. The tail binds to the head-neck domain, inhibiting ATPase activity of myosin VIIA. Proc Natl Acad Sci U S A 106, 8483–8488 (2009).

6. Umeki, N. et al. Phospholipid-dependent regulation of the motor activity of myosin X. Nat Struct Mol Biol 18, 783–788 (2011).

7. Spink, B. J., Sivaramakrishnan, S., Lipfert, J., Doniach, S. & Spudich, J. A. Long single alpha-helical tail domains bridge the gap between structure and function of myosin VI. Nat Struct Mol Biol 15, 591–597 (2008).

8. Lister, I. et al. A monomeric myosin VI with a large working stroke. EMBO J 23, 1729–1738 (2004).

9. Caporizzo, M. A. et al. The Antiparallel Dimerization of Myosin X Imparts Bundle Selectivity for Processive Motility. Biophys J 114, 1400–1410 (2018).

10. Liu, R. et al. A binding protein regulates myosin-7a dimerization and actin bundle assembly. Nat Commun 12, 563 (2021).

11. Lu, Q., Ye, F., Wei, Z., Wen, Z. & Zhang, M. Antiparallel coiled-coil-mediated dimerization of myosin X. Proc Natl Acad Sci U S A 109, 17388–17393 (2012).

12. Park, H. et al. Full-length myosin VI dimerizes and moves processively along actin filaments upon monomer clustering. Mol Cell 21, 331–336 (2006).

13. Ropars, V. et al. The myosin X motor is optimized for movement on actin bundles. Nat Commun 7, 12456 (2016).

14. Yu, C. et al. Myosin VI undergoes cargo-mediated dimerization. Cell 138, 537–548 (2009).

15. Phichith, D. et al. Cargo binding induces dimerization of myosin VI. Proc Natl Acad Sci U S A 106, 17320–17324 (2009).

16. Kim, H., Hsin, J., Liu, Y., Selvin, P. R. & Schulten, K. Formation of salt bridges mediates internal dimerization of myosin VI medial tail domain. Structure 18, 1443–1449 (2010).

17. Mukherjea, M. et al. Myosin VI must dimerize and deploy its unusual lever arm in order to perform its cellular roles. Cell Rep 8, 1522–1532 (2014).

18. Mukherjea, M. et al. Myosin VI dimerization triggers an unfolding of a three-helix bundle in order to extend its reach. Mol Cell 35, 305–315 (2009).

19. de Jonge, J. J., Batters, C., O’Loughlin, T., Arden, S. D. & Buss, F. The MYO6 interactome: selective motor-cargo complexes for diverse cellular processes. FEBS Lett 593, 1494–1507 (2019).

20. Avraham, K. B. et al. The mouse Snell’s waltzer deafness gene encodes an unconventional myosin required for structural integrity of inner ear hair cells. Nat Genet 11, 369–375 (1995).

21. Ripoll, L. et al. Myosin VI and branched actin filaments mediate membrane constriction and fission of melanosomal tubule carriers. J Cell Biol 217, 2709–2726 (2018).

22. Tumbarello, D. A. et al. Autophagy receptors link myosin VI to autophagosomes to mediate Tom1-dependent autophagosome maturation and fusion with the lysosome. Nat Cell Biol 14, 1024–1035 (2012).

23. Morris, S. M. et al. Myosin VI binds to and localises with Dab2, potentially linking receptor-mediated endocytosis and the actin cytoskeleton. Traffic 3, 331–341 (2002).

24. Naccache, S. N., Hasson, T. & Horowitz, A. Binding of internalized receptors to the PDZ domain of GIPC/synectin recruits myosin VI to endocytic vesicles. Proc Natl Acad Sci U S A 103, 12735–12740 (2006).

25. Fili, N. et al. NDP52 activates nuclear myosin VI to enhance RNA polymerase II transcription. Nat Commun 8, 1871 (2017).

26. Yoshida, H. et al. Lessons from border cell migration in the Drosophila ovary: A role for myosin VI in dissemination of human ovarian cancer. Proc Natl Acad Sci U S A 101, 8144–8149 (2004).

27. Wang, D. et al. MYO6 knockdown inhibits the growth and induces the apoptosis of prostate cancer cells by decreasing the phosphorylation of ERK1/2 and PRAS40. Oncol Rep 36, 1285–1292 (2016).

28. Fili, N. et al. Competition between two high- and low-affinity protein-binding sites in myosin VI controls its cellular function. J Biol Chem 295, 337–347 (2020).

29. Dos Santos, Á. et al. Binding partners regulate unfolding of myosin VI to activate the molecular motor. Biochem J 479, 1409–1428 (2022).

30. Hariadi, R. F., Cale, M. & Sivaramakrishnan, S. Myosin lever arm directs collective motion on cellular actin network. Proceedings of the National Academy of Sciences 111, 4091–4096 (2014).

31. Rai, A., Vang, D., Ritt, M. & Sivaramakrishnan, S. Dynamic multimerization of Dab2-Myosin VI complexes regulates cargo processivity while minimizing cortical actin reorganization. J Biol Chem 296, 100232 (2021).

32. Altman, D., Sweeney, H. L. & Spudich, J. A. The mechanism of myosin VI translocation and its load-induced anchoring. Cell 116, 737–749 (2004).

33. Rai, A. et al. Multimodal regulation of myosin VI ensemble transport by cargo adaptor protein GIPC. J Biol Chem 298, 101688 (2022).

34. Shang, G. et al. Structure analyses reveal a regulated oligomerization mechanism of the PlexinD1/GIPC/myosin VI complex. Elife 6, e27322 (2017).

35. Hu, S. et al. Structure of Myosin VI/Tom1 complex reveals a cargo recognition mode of Myosin VI for tethering. Nat Commun 10, 3459 (2019).

36. Batters, C., Brack, D., Ellrich, H., Averbeck, B. & Veigel, C. Calcium can mobilize and activate myosin-VI. Proc Natl Acad Sci U S A 113, E1162–1169 (2016).

37. Bonnet, J. et al. Autocatalytic association of proteins by covalent bond formation: a Bio Molecular Welding toolbox derived from a bacterial adhesin. Sci Rep 7, 43564 (2017).

38. Brownstein, Z. et al. Novel myosin mutations for hereditary hearing loss revealed by targeted genomic capture and massively parallel sequencing. Eur J Hum Genet 22, 768–775 (2014).

39. Dennis, M. K. et al. BLOC-1 and BLOC-3 regulate VAMP7 cycling to and from melanosomes via distinct tubular transport carriers. J Cell Biol 214, 293–308 (2016).

40. Ishida, M., Arai, S. P., Ohbayashi, N. & Fukuda, M. The GTPase-deficient Rab27A(Q78L) mutant inhibits melanosome transport in melanocytes through trapping of Rab27A effector protein Slac2-a/melanophilin in their cytosol: development of a novel melanosome-targetinG tag. J Biol Chem 289, 11059–11067 (2014).

41. He, F. et al. Myosin VI Contains a Compact Structural Motif that Binds to Ubiquitin Chains. Cell Rep 14, 2683–2694 (2016).

42. Berggren, K. et al. Background-free, high sensitivity staining of proteins in one- and two-dimensional sodium dodecyl sulfate-polyacrylamide gels using a luminescent ruthenium complex. Electrophoresis 21, 2509–2521 (2000).

43. Yu, C. et al. Membrane-induced lever arm expansion allows myosin VI to walk with large and variable step sizes. J Biol Chem 287, 35021–35035 (2012).

44. Rock, R. S. et al. Myosin VI is a processive motor with a large step size. Proceedings of the National Academy of Sciences 98, 13655–13659 (2001).

45. Spitzer, M., Wildenhain, J., Rappsilber, J. & Tyers, M. BoxPlotR: a web tool for generation of box plots. Nat Methods 11, 121–122 (2014).

46. Sweeney, H. L. & Houdusse, A. What can myosin VI do in cells? Curr Opin Cell Biol 19, 57–66 (2007).

47. Lumb, K. J., Carr, C. M. & Kim, P. S. Subdomain folding of the coiled coil leucine zipper from the bZIP transcriptional activator GCN4. Biochemistry 33, 7361–7367 (1994).

48. Heissler, S. M. & Sellers, J. R. Various Themes of Myosin Regulation. J Mol Biol 428, 1927–1946 (2016).

49. Fili, N. & Toseland, C. P. Unconventional Myosins: How Regulation Meets Function. Int J Mol Sci 21, E67 (2019).

50. Jumper, J. et al. Highly accurate protein structure prediction with AlphaFold. Nature 596, 583–589 (2021).

51. Varadi, M. et al. AlphaFold Protein Structure Database: massively expanding the structural coverage of protein-sequence space with high-accuracy models. Nucleic Acids Res 50, D439–D444 (2022).

52. Spudich, G. et al. Myosin VI targeting to clathrin-coated structures and dimerization is mediated by binding to Disabled-2 and PtdIns(4,5)P2. Nat Cell Biol 9, 176–183 (2007).

53. Thureau, A., Roblin, P. & Pérez, J. BioSAXS on the SWING beamline at Synchrotron SOLEIL. J Appl Cryst 54, 1698–1710 (2021).

54. Manalastas-Cantos, K. et al. ATSAS 3.0: expanded functionality and new tools for small-angle scattering data analysis. J Appl Crystallogr 54, 343–355 (2021).

55. Svergun, D. I., Petoukhov, M. V. & Koch, M. H. Determination of domain structure of proteins from X-ray solution scattering. Biophys J 80, 2946–2953 (2001).

56. Volkov, V. V., Svergun, D. I. & IUCr. Uniqueness of ab initio shape determination in small-angle scattering. Journal of Applied Crystallography vol. 36 860–864 http://scripts.iucr.org/cgi-bin/paper?S0021889803000268 (2003).

57. De La Cruz, E. M., Ostap, E. M. & Sweeney, H. L. Kinetic mechanism and regulation of myosin VI. J Biol Chem 276, 32373–32381 (2001).

58. Punjani, A., Rubinstein, J. L., Fleet, D. J. & Brubaker, M. A. cryoSPARC: algorithms for rapid unsupervised cryo-EM structure determination. Nat Methods 14, 290–296 (2017).

59. Ménétrey, J. et al. Processive steps in the reverse direction require uncoupling of the lead head lever arm of myosin VI. Mol Cell 48, 75–86 (2012).

60. Barnes, C. A. et al. Remarkable Rigidity of the Single α-Helical Domain of Myosin-VI As Revealed by NMR Spectroscopy. J Am Chem Soc 141, 9004–9017 (2019).

61. Schrödinger, L. & DeLano, W. PyMOL. Retrieved from http://www.pymol.org/pymol. (2020).

62. Roth, S. & Heintzmann, R. Optical photon reassignment with increased axial resolution by structured illumination. Methods Appl Fluoresc 4, 045005 (2016).

63. Vonrhein, C. et al. Data processing and analysis with the autoPROC toolbox. Acta Crystallogr D Biol Crystallogr 67, 293–302 (2011).

64. Tickle, I. J. et al. STARANISO. Cambridge, United Kingdom: Global Phasing Ltd. (2018).

65. McCoy, A. J. et al. Phaser crystallographic software. J Appl Cryst 40, 658–674 (2007).

66. Terwilliger, T. C. et al. Iterative model building, structure refinement and density modification with the PHENIX AutoBuild wizard. Acta Crystallogr D Biol Crystallogr 64, 61–69 (2008).

67. Emsley, P. & Cowtan, K. Coot: model-building tools for molecular graphics. Acta Crystallogr D Biol Crystallogr 60, 2126–2132 (2004).

68. Bricogne, G. et al. BUSTER version X.Y.Z. Cambridge, United Kingdom: Global Phasing Ltd. (2017).

69. Gurel, P. S. et al. Cryo-EM structures reveal specialization at the myosin VI-actin interface and a mechanism of force sensitivity. Elife 6, e31125 (2017).

70. Berman, H., Henrick, K. & Nakamura, H. Announcing the worldwide Protein Data Bank. Nat Struct Biol 10, 980 (2003).

