## Supplementary informations for "How Myosin VI Traps its Off-State, is Activated and Dimerizes"

### Supplementary Figures

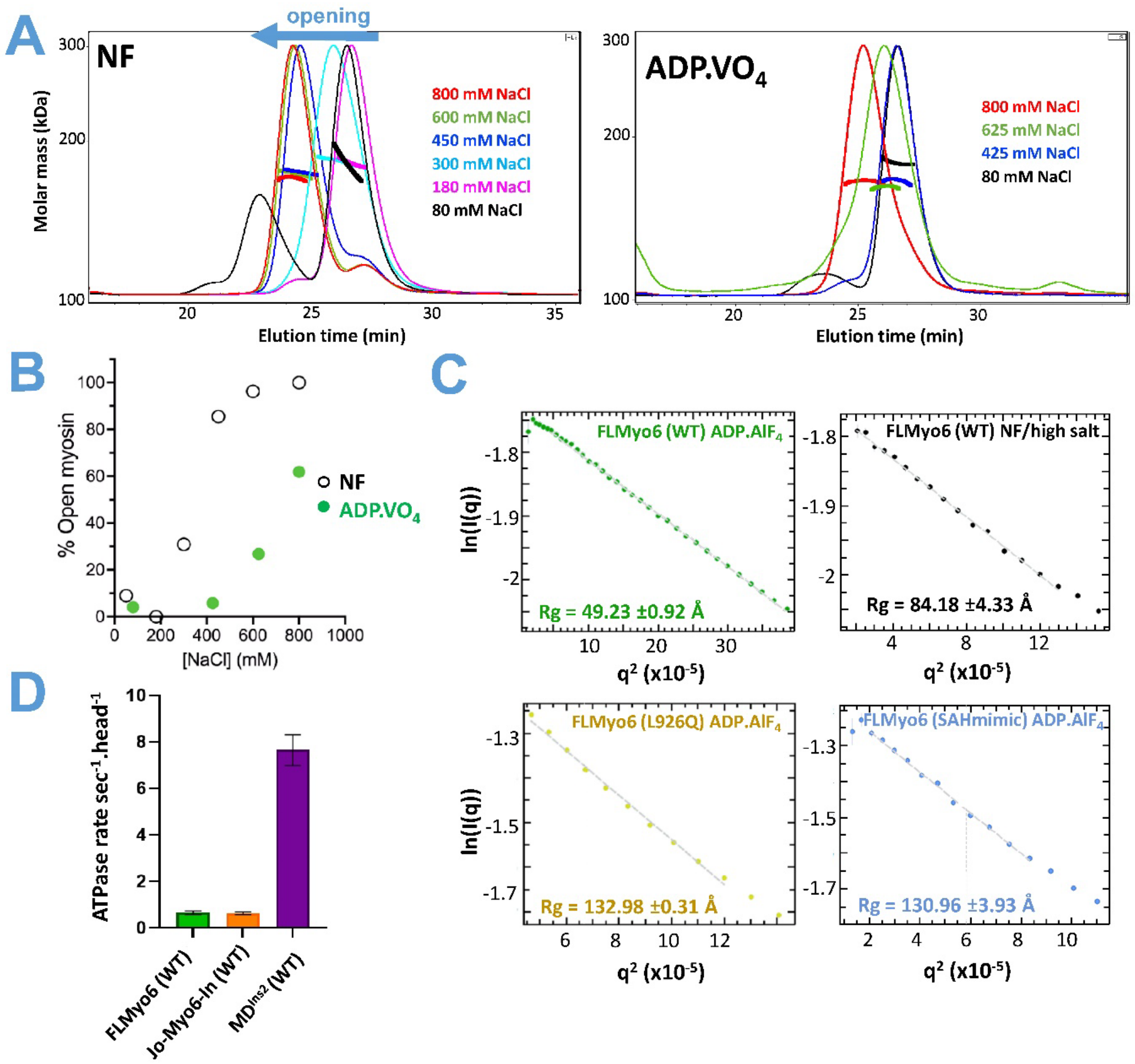


***Supplementary Figure 1 – ATP is required for the compact, back-folded conformation of Myo6.***

**(A)** Normalized SEC-MALS profiles of FLMyo6 in different elution buffers. Thin lines: normalized static light scattering; Thick lines: molecular mass as determined for each 0.5 second experimental point by Astra (Wyatt technology). ADP.$VO_4$ buffer: 20 mM Hepes; 2 mM $MgCl_2$; 1 mM NaADP; 1 mM $NaVO_4$; 0.1 mM EGTA; 0.5 mM TCEP, pH 7.5 + indicated NaCl concentration; NF buffer: 20 mM Hepes; 2 mM $MgCl_2$; 0.1 mM EGTA; 0.5 mM TCEP, pH 7.5 + indicated NaCl concentration.

**(B)** Percentage of open FLMyo6 as a function of [NaCl] in ADP.$VO_4$ or NF conditions, as determined by SEC-MALS. The earliest and latest elution times for Myo6 monomers were taken as 100% and 0% open, respectively.

**(C)** Guinier plot of FLMyo6 WT, SAHmimic and L926Q in ADP.$AlF_4$ or in NF/high salt. Rg values were extracted from the linear fit (dashed line) using primusqt (ATSAS suite[1]).

**(D)** Actin-activated ATPase rate of 150 nM FLMyo6 (WT), Jo-Myo6-In, and $MD^{Ins2}$ (n=6) at 40 µM F-actin.

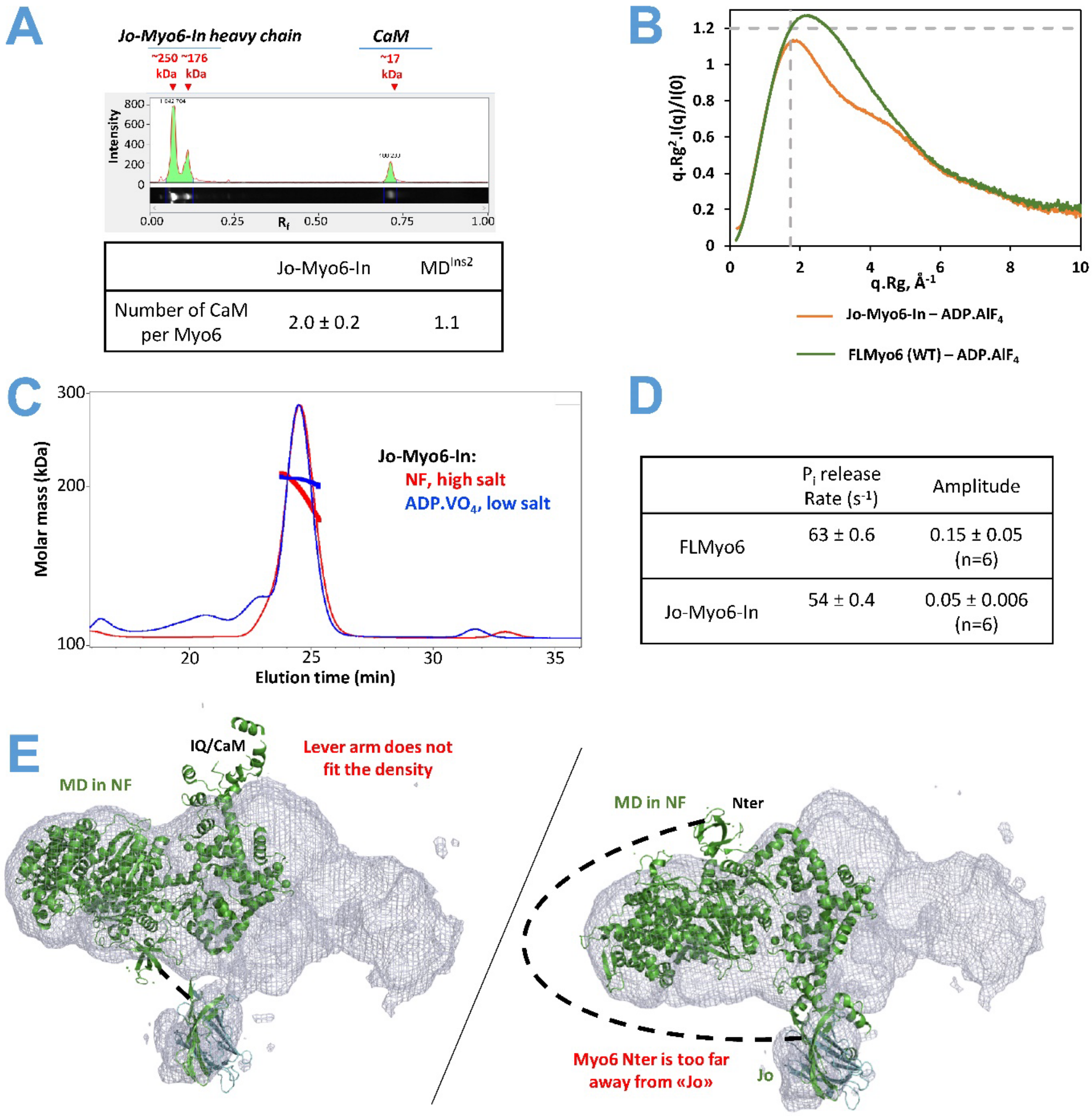


***Supplementary Figure 2 – The Jo-In fusion stabilizes the off-state with no effect on its conformation.*** **(A)** SDS-PAGE profile of Jo-Myo6-In stained with SYPRO[2]. The estimated molecular weight based on SDS-PAGE is given in red. The CaM/FLMyo6 molar ratio was determined from the intensities of the corresponding bands (Table: mean ± standard deviation from 4 replicates). Intensity of Myo6 and CaM bands were estimated on ImageLab software (Bio-Rad). Ratio CaM/FLMyo6 was estimated by calculating (CaM band intensity)/(Myo6 band intensity) and normalized by theoretical molecular weight to get a molar ratio. $MD^{Ins2}$ was used as a control as it binds 1 CaM per Myo6[3]. Note that, on SDS-PAGE, purified Jo-Myo6-In-Flag appears as two distinct bands, one at its expected molecular weight 176 kDa, and another band around 250 kDa . On SEC, the faster band was found to elute slightly earlier, in a broader peak merged to the one corresponding to the slower band. LC-MS concluded that both bands correspond to Jo-Myo6-In with 68.5% sequence coverage. Exactly the same peptides were found in both bands, and the Jo and In peptides containing the expected covalent Lys-Asn bond could not be found. Therefore, we assume that the slower band corresponds to the protein whose covalent Lys-Asn bond is correctly formed (=”locked”), whereas the faster band is composed by folded, but not locked, protein (=”closed”).

**(B)** Dimensionless Kratky plot representation. Dotted lines ($(qRg)^2I(q)/I(0) = 1.104$ and $qRg = \sqrt{3}$) intersection highlight the theoretical maximum of a globular protein. In the presence of ADP.$AlF_4$, FLMyo6 (green) and Jo-Myo6-In (orange) spectra result in a bell-shape curve (with a shoulder in the case of Jo-In) with a maximum close to ($\sqrt{3}$:1.104), suggesting that both proteins are rather globular and well folded. The shoulder in the Jo-Myo6-In profile is typical of multi-domain proteins consistent with the addition of the Jo-In tag.
**(C)** Normalized SEC-MALS profiles of Jo-Myo6-In in conditions that favor opening (red, NF) or closure (blue, ADPVO4). NF buffer: 20 mM Hepes; 300 mM NaCl; 2 mM $CaCl_2$; 0.1 mM EDTA; 0.5 mM TCEP, pH 7.5. ADP.$VO_4$ buffer: 20 mM Hepes; 80 mM NaCl; 2 mM $MgCl_2$; 1 mM NaADP; 1 mM $NaVO_4$; 0.1 mM EGTA; 0.5 mM TCEP, pH 7.5.
**(D)** Myo6 $P_i$ release rates and amplitudes from stopped-flow experiments.
**(E)** Docking of nucleotide-free Myo6 $MD^{Ins2/IQ}$ (PDB: 2BKI[3]) and Jo-In (PDB: 5MKC[4]) in the negative staining map. The structure of the Motor domain devoid of nucleotide doesn't fit the map: **(left)** if the connection between the N-terminus of Myo6 and the C-terminus of Jo is respected, the Lever arm does not fit in the envelope; **(right)** if the Lever arm is positioned inside the 3D reconstruction, the distance between the Myo6 N-terminus and the fusion subdomain Jo is prohibitive.

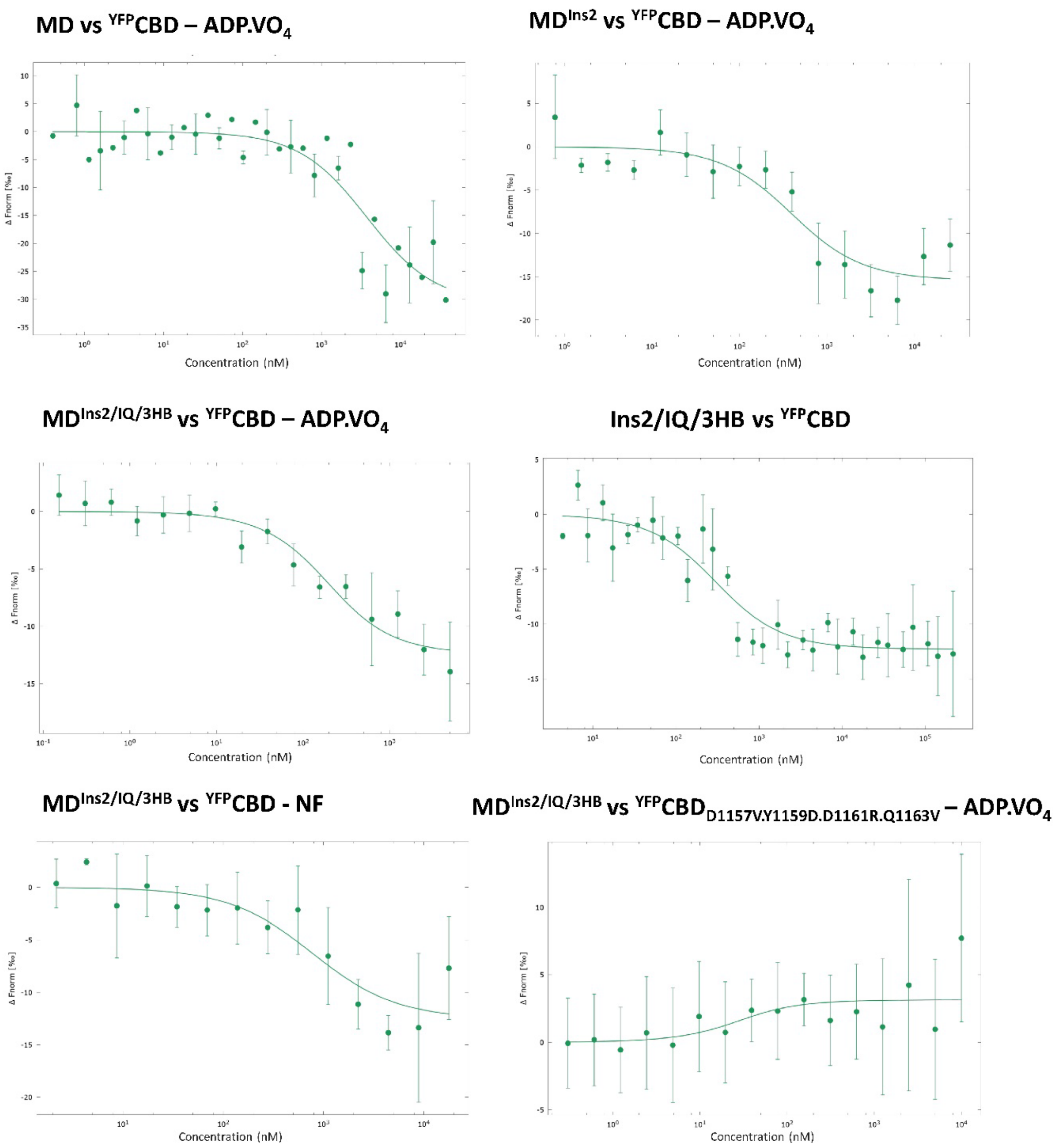


***Supplementary Figure 3 – Affinity of the Myo6 CBD for different Myo6 Head or Neck constructs.***

Microscale thermophoresis profiles and fits corresponding to the data exposed in Table 1.

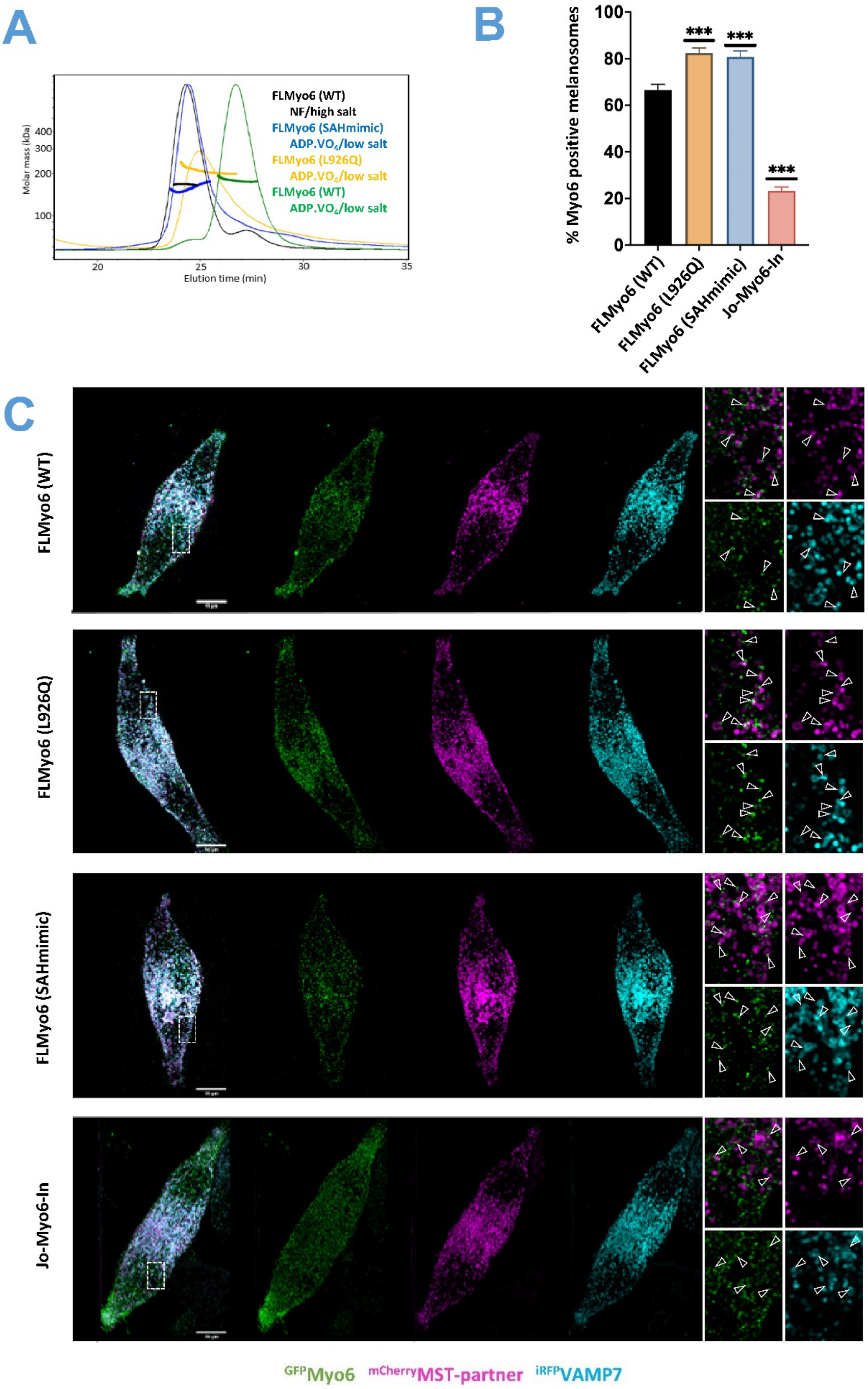


***<u>Supplementary Figure 4 – Role of the proximal Myo6 sequence in the stabilization of the off-state.</u>***
**(A)** Normalized SEC-MALS profiles of FLMyo6 (SAHmimic) (blue) and FLMyo6 (L926Q) (yellow) mutants, plus FLMyo6 (WT) in conditions that favor opening (black) or closure (green), as references.

Experimental conditions were similar to those described for Sup Fig 1. Here, “High salt” means 800 mM NaCl; “low salt”, 80 mM.

**(B)** Quantification of Myo6-positive melanosomes for $^{GFP}$FLMyo6 (L926Q), $^{GFP}$FLMyo6 (SAHmimic), $^{GFP}$Jo-Myo6-In, and $^{GFP}$FLMyo6 (WT) (n=3, total cell number ~30). Significance: ***, P < 0.001 (unpaired t test with Welch’s correction).

**(C)** Representative fixed MNT-1 cells co-expressing $^{GFP}$FLMyo6 (WT), $^{GFP}$Jo-Myo6-In, $^{GFP}$FLMyo6 (SAHmimic) or $^{GFP}$FLMyo6 (L926Q) with $^{mCherry}$MST and $^{iRFP}$VAMP7. $^{mCherry}$MST and $^{iRFP}$VAMP7 are melanosome-associated components. MNT-1 cells were fixed 48h post-transfection, then imaged and processed for quantification. Green: $^{GFP}$Myo6; Magenta: $^{mCherry}$MST; Cyan: $^{iRFP}$VAMP7. From left to right: Entire cell: 3 channels merge and individuals; 8x zoom of the boxed region: merged $^{GFP}$Myo6 / $^{mCherry}$MST, individual channels. Scale bars: 10 µm.

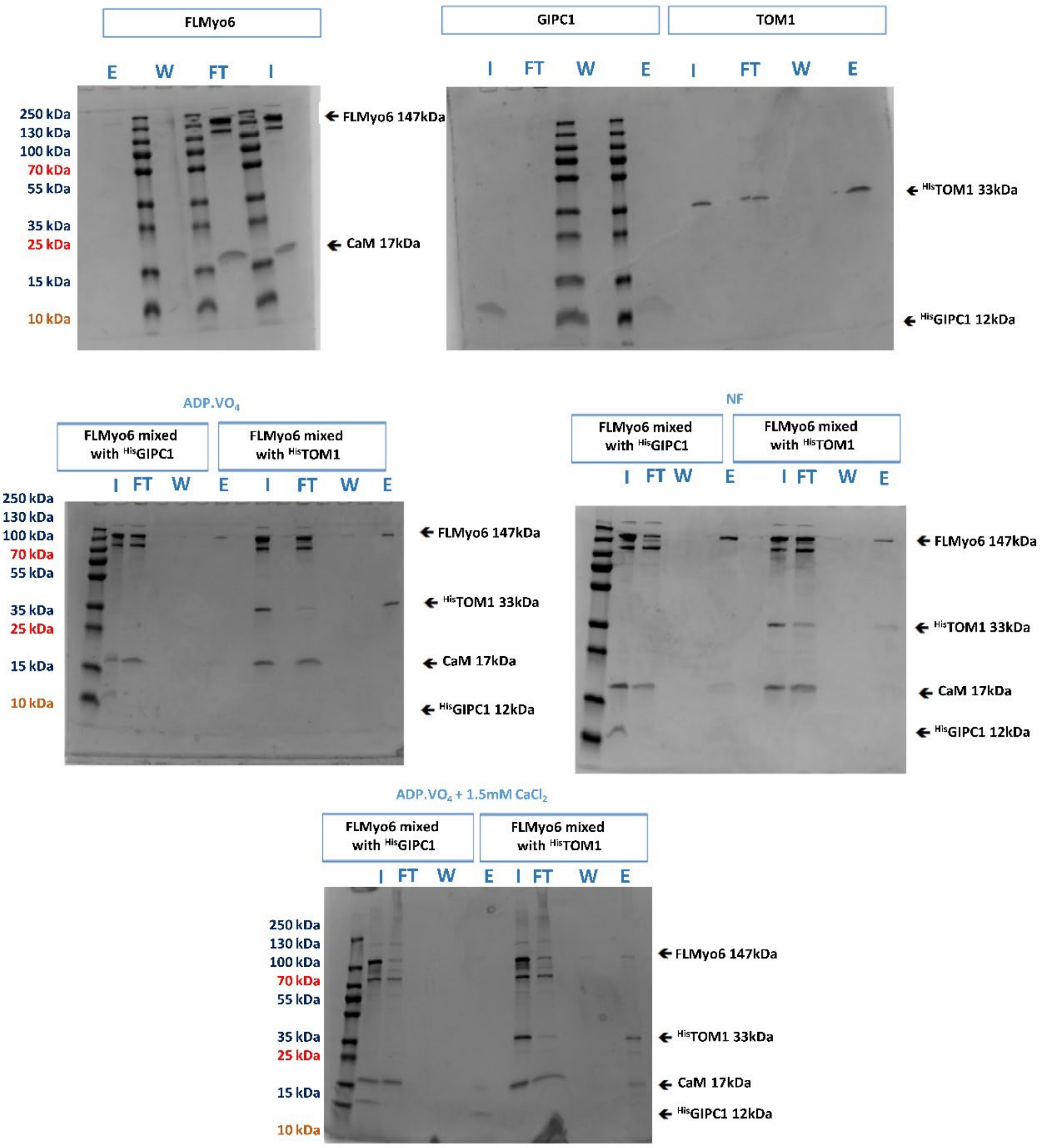


***Supplementary Figure 5 – SDS-PAGE stained with colloidal blue corresponding to Anti-His pull-down in Fig. 3D.***

Input (I), flow through (FT), last wash (W) and elution (E) fractions corresponding to anti-His pull-down pictured in Fig. 3D.

| | Partner | Myo6 Tail | $K_D$ | n |
|---|---|---|---|---|
| **Interaction with GIPC1** | $^{His}$GIPC1 | $^{YFP}$CBD ( WT) | 214 ± 144 nM | 2 |
| | $^{His}$GIPC1 | $^{YFP}$CBD (I1072A) | 227 ± 108 nM | 2 |
| **Interaction with Dab2** | $^{His}$Dab2 | $^{YFP}$CBD ( WT) | 423 ± 346 nM | 2 |
| **Interaction with TOM1** | $^{His}$TOM1 | $^{YFP}$CBD ( WT) | 445 ± 135 nM | 2 |

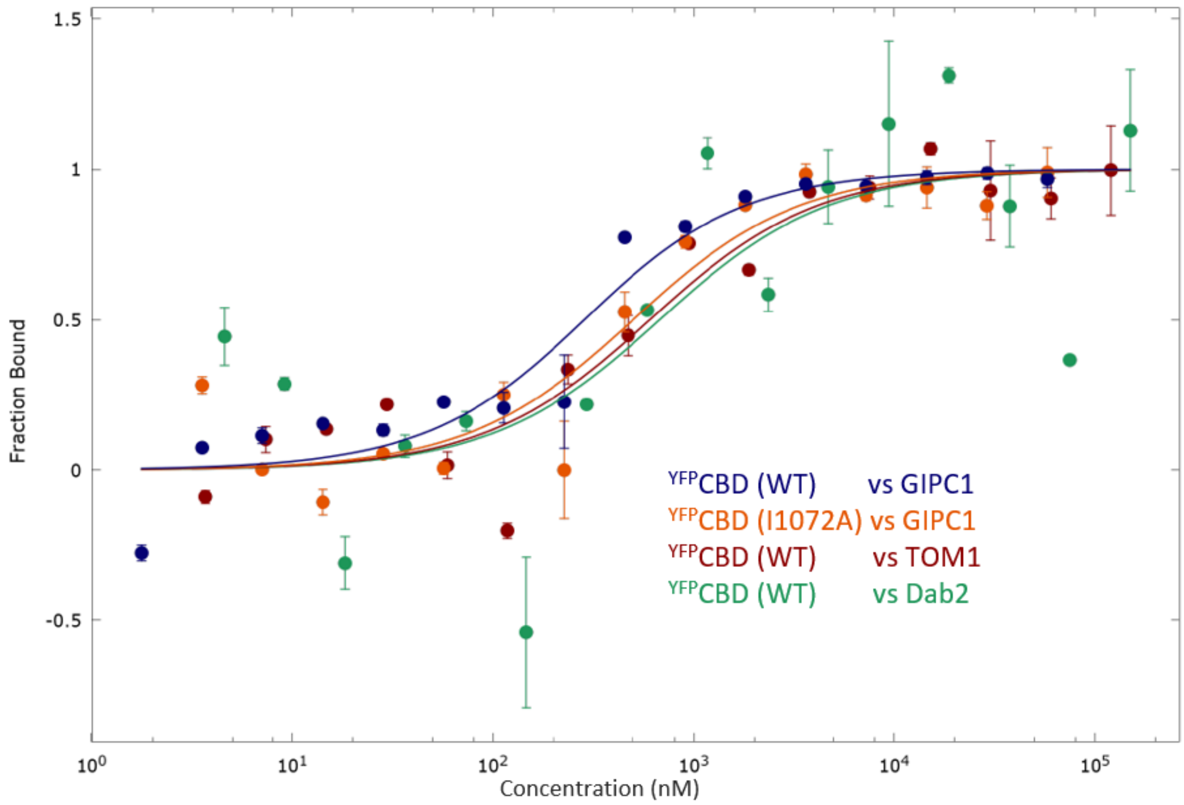


***<u>Supplementary Figure 6 – Affinity of the Myo6 tail for partners by Microscale Thermophoresis.</u>***

**(Top)** Dissociation constant ($K_D$) ± $K_D$ confidence determined by microscale thermophoresis of Myo6 CBD constructs against Myo6 partners. **(Bottom)** Integration of thermophoresis profiles against concentration of partners (nM) for one replicate shown as an example. The affinity was quantified by analyzing the change in thermophoresis as a function of the concentration of the titrated protein using the NTAnalysis software provided by the manufacturer. Error bars: standard deviation. Fit: “Kd model” equation from NTAnalysis.

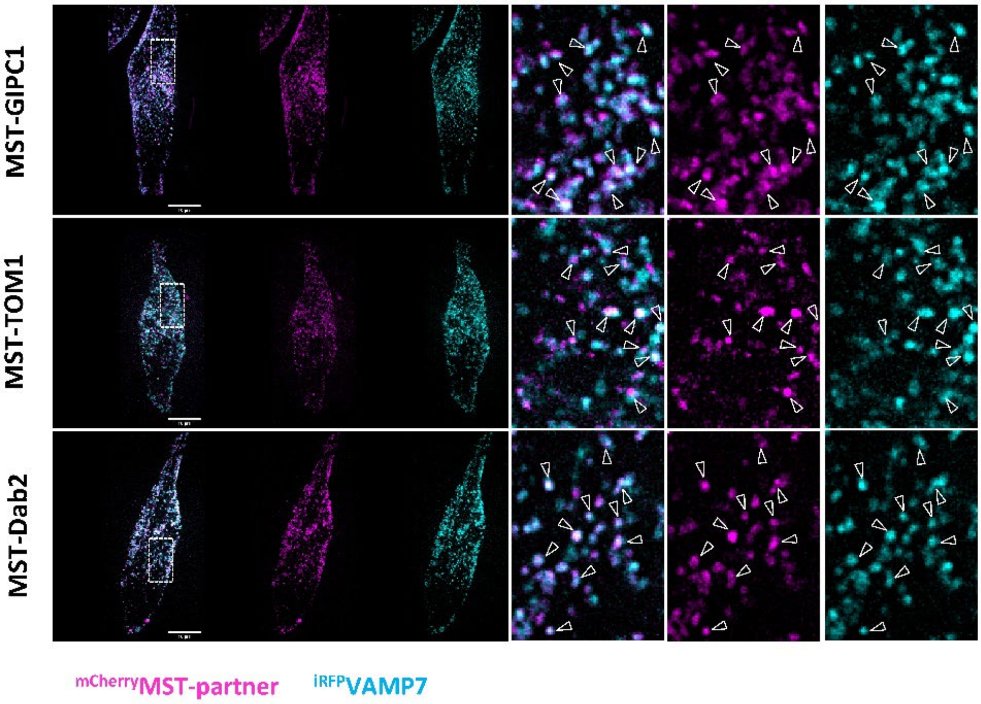


***Supplementary Figure 7 – Localization of MST-partners at melanosomal membrane in MNT-1 cells.*** Representative MNT-1 cells co-expressing $^{mCherry}$MST-partners and $^{iRFP}$VAMP7. MNT-1 were fixed 48h post-transfection then imaged. In order: $^{mCherry}$MST-GIPC1, $^{mCherry}$MST-TOM1, $^{mCherry}$MST-Dab2. Magenta: $^{mCherry}$MST-partners; Cyan: $^{iRFP}$VAMP7. From left to right: entire cell (3 channels merged, individual channels); zoom 8x boxed region ($^{mCherry}$MST-partners / $^{iRFP}$VAMP7 merged, individual channels). Results show colocalization of $^{mCherry}$MST-partners all around the $^{iRFP}$VAMP7 melanosomes.

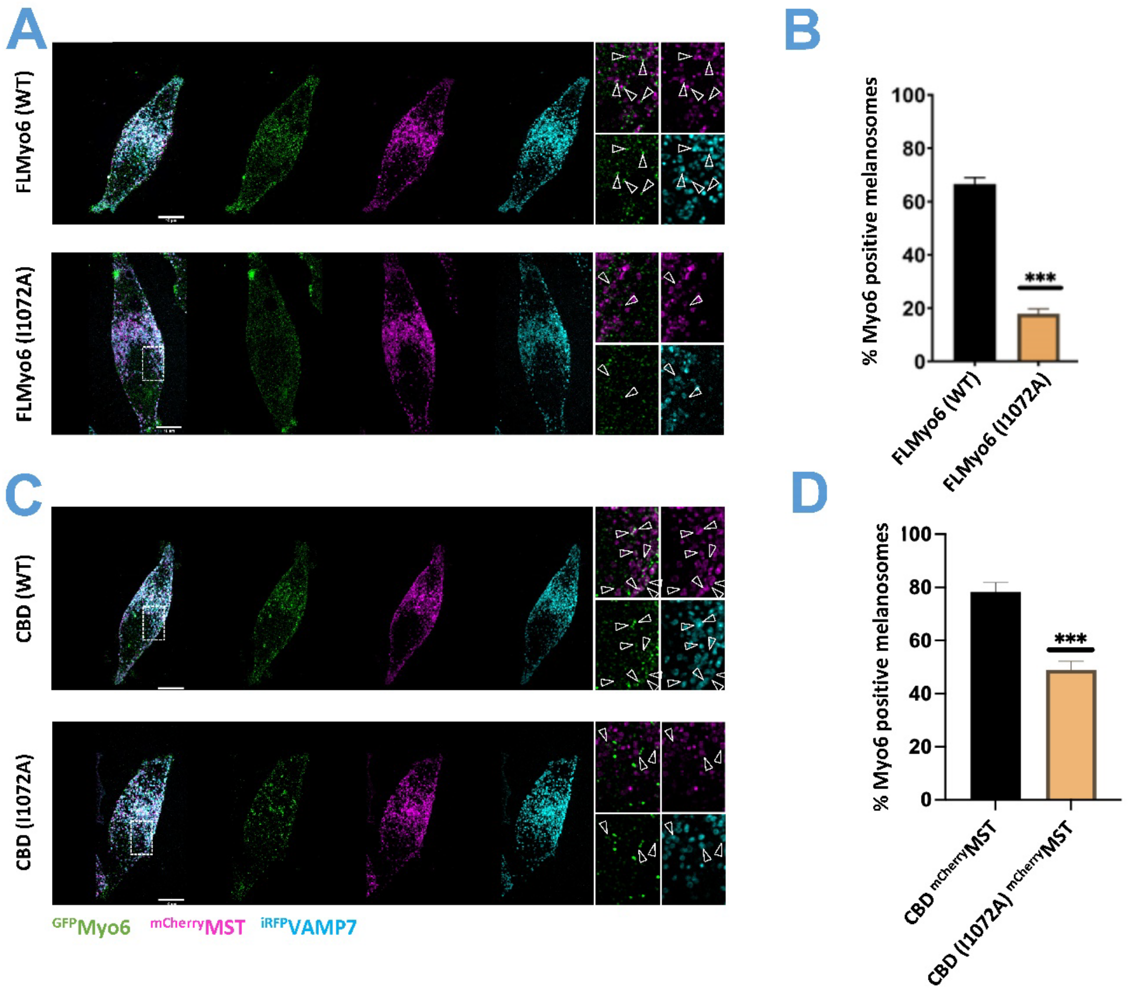


***<u>Supplementary Figure 8 – I1072 is key for Myo6 specific recruitment on melanosomes.</u>***

**(A)** Representative fixed MNT-1 cells co-expressing $^{GFP}$Myo6 +/- I1072A mutation with $^{mcherry}$MST and $^{iRFP}$VAMP7. $^{mCherry}$MST and $^{iRFP}$VAMP7 are melanosome-associated components. MNT-1 cells were fixed 48h post-transfection then imaged and processed for quantification. Note that the control FLMyo6 (WT) is the same as in Sup Fig. 4B.

**(B)** Quantification of Myo6-positive melanosomes for $^{GFP}$FLMyo6 (WT) and $^{GFP}$FLMyo6 (I1072A) in MNT-1 cells (n=3, total cell number ~30).

**(C)** Representative MNT-1 cells co-expressing Myo6 CBD +/- I1072A mutation with $^{mcherry}$MST and $^{iRFP}$VAMP7. $^{mCherry}$MST and $^{iRFP}$VAMP7 are melanosome-associated components. MNT-1 cells were fixed 48h post-transfection then imaged and processed for quantification.

**(D)** Quantification of Myo6-positive melanosomes for $^{GFP}$CBD (WT) and $^{GFP}$CBD (I1072A) in MNT-1 cells (n=2, total cell number ~20).

**(A, C)** Green: $^{GFP}$Myo6 constructs; Cyan: $^{iRFP}$VAMP7; Magenta: $^{mcherry}$MST-partner. Entire cell merge 3 channels and individuals. Zoom (8x boxed region): merge $^{GFP}$Myo6 / $^{mcherry}$MST-partner, individual channels. Scale bars: 10µm.

**(C, D)** *Myo6-positive melanosomes are expressed in percentage and normalized to the total number of VAMP7-positive melanosomes.* Significance: ***, $P < 0.001$ (unpaired t test with Welch's correction), *for each $^{GFP}$Myo6 construct, significance of experiments with partners compared to the control without partner (in black on the graph).*

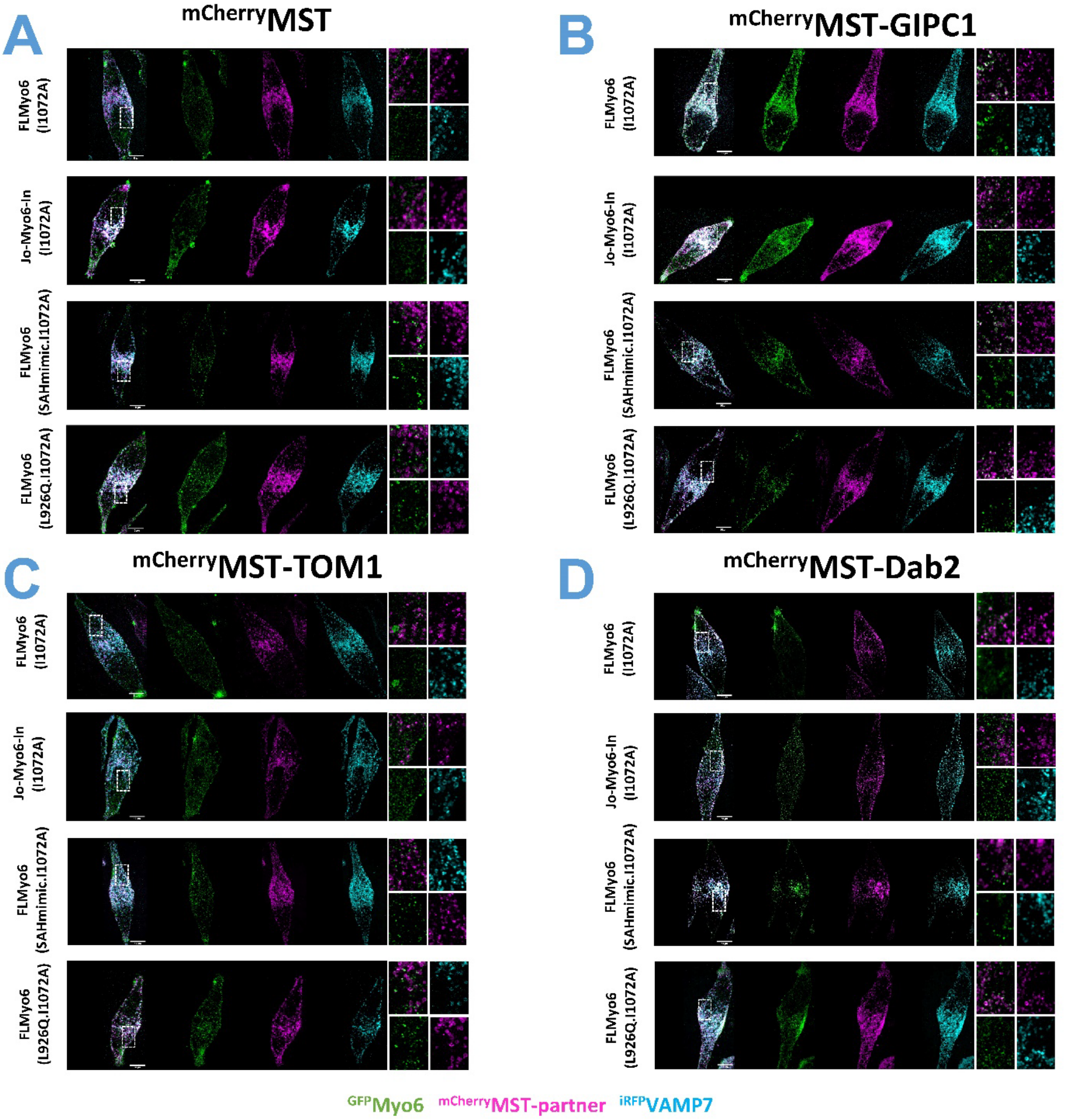


***<u>Supplementary Figure 9 – GIPC1 can bind the back-folded Myo6 state and activate it, while Dab2 and Tom1 can only bind to Myo6 once the motor has been primed open.</u>***

Representative entire MNT-1 cells corresponding to the zooms in Fig. 4. Left to right: merge of 3 channels, individual channels. Zoom (8x boxed region): merge $^{GFP}$Myo6 / $^{mCherry}$MST-partners, individual channels. Scale bars: 10µm. $^{GFP}$FLMyo6 constructs: $^{GFP}$FLMyo6 (I1072A), $^{GFP}$FLMyo6 (SAHmimic.I1072A), $^{GFP}$FLMyo6 (L926Q.I1072A), Jo-Myo6-In (I1072A).

**(A)** Representative fixed MNT-1 cells co-expressing different $^{GFP}$FLMyo6 (I1072A) constructs with control $^{mCherry}$MST and $^{iRFP}$VAMP7.

**(B)** Representative fixed MNT-1 cells co-expressing different $^{GFP}$FLMyo6 (I1072A) constructs with $^{mCherry}$MST-GIPC1 and $^{iRFP}$VAMP7.

**(C)** Representative fixed MNT-1 cells co-expressing different $^{GFP}$FLMyo6 (I1072A) constructs with $^{mCherry}$MST-TOM1 and $^{iRFP}$VAMP7.

**(D)** Representative fixed MNT-1 cells co-expressing different $^{GFP}$FLMyo6 (I1072A) constructs with $^{mCherry}$MST-Dab2 and $^{iRFP}$VAMP7.

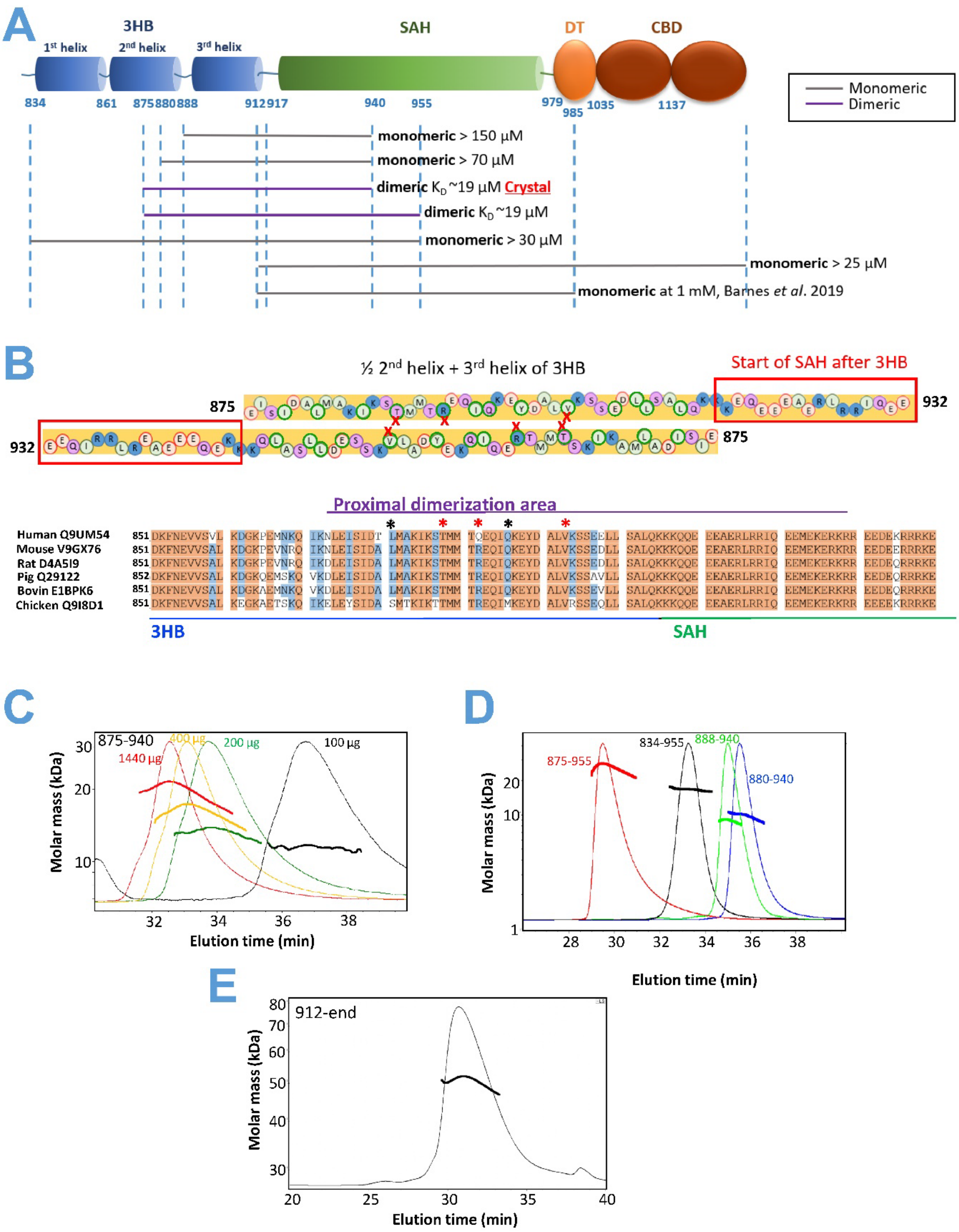


***__Supplementary Figure 10 –Characterization of the dimerization of proximal region by MALS.__***

The 875-940 fragment is sufficient for dimerization. Importantly, a peptide missing the first 5 aa (aa 880-940) failed to dimerize even when its peak concentration was higher than 70 µM, which emphasised the key role of residues 875-880 for the stability of the dimerization. Additionally, in line with a recent study[5], no dimerization was found via the SAH, unlike previously proposed by molecular dynamics simulations[6]. Indeed, no dimerization was observed for the Tail fragment aa 912-end, (containing the whole SAH), even when its peak concentration was 25 µM.

(A) Scheme of the Myo6 Tail showing the oligomerization states found for different constructs using SEC-MALS in 10 mM Tris pH 7.5; 50 mM NaCl; 5 mM $NaN_3$; 0.5 mM TCEP.

(B) (Top) schematic representation of the dimer crystal structure. Negatively charged residues are circled in red, positively charged residues are circled in blue, hydrophobic residues are circled in green and polar, uncharged residues are circled in purple. Residues involved in apolar contacts are circled in green. Three residues found to be essential for dimerization (T888D, R892E, V903D) are marked with a red cross. (Bottom) Sequence alignment of 851-949 (human protein numbering). Red: conserved residues; blue: conserved in 5 out of 6 species. *: residues inside the dimerization area that are not 100% conserved (S881 and M896). In the structure, M896 faces itself and seems compatible with the crystallized dimer. *: residues mutated in this study to disrupt proximal dimerization. Residues T888 and V903 are well conserved. The crystal structure shows that the replacement of R892 by Q in the human sequence should be compatible with proximal dimerization.

(C) Titration of the 875-940 fragment by SEC-MALS in 10 mM Tris pH 7.5; 50 mM NaCl; 5 mM $NaN_3$; 0.5 mM TCEP. Thin lines: light scattering (normalized); thick lines: molecular masses. A $K_D^{App}$ of ~19 µM was calculated from a fit of molecular weight against the concentration at the peak from each profile – the affinity is probably underestimated because of the size exclusion resin.

(D) Normalized SEC-MALS profiles of the 834-955, 880-940 and 888-940 constructs (concentration at the peak stated on Sup Fig. 10A) found to be monomeric in 10 mM Tris pH 7.5; 50 mM NaCl; 5 mM $NaN_3$; 0.5 mM TCEP, unlike the dimeric 875-955 construct (red).

(E) Normalized SEC-MALS profile of the 912-end construct (25 µM at the peak) found to be monomeric in 10 mM Tris pH 7.5; 50 mM NaCl; 5 mM $NaN_3$; 0.5 mM TCEP.

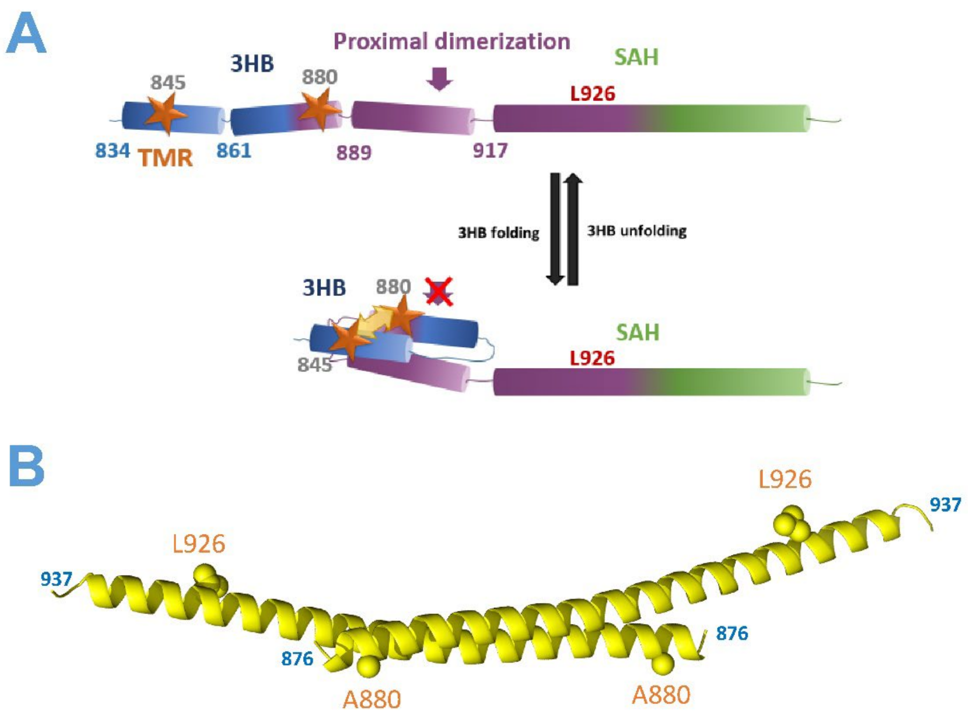


***<u>Supplementary Figure 11 – Proximal dimerization is regulated by 3HB unfolding which is impaired by the pathogenic L926Q deafness mutation.</u>***

**(A)** Scheme representing Myo6 SAH (green) and 3HB (blue), with the proximal dimerization area (875-940) shown in purple. Stars: TMR fluorescent spots. Their proximity promotes quenching when the 3HB is folded.

**(B)** Crystal structure of the proximal dimer pictured in cartoon, residues A880 and L926 are shown as spheres.

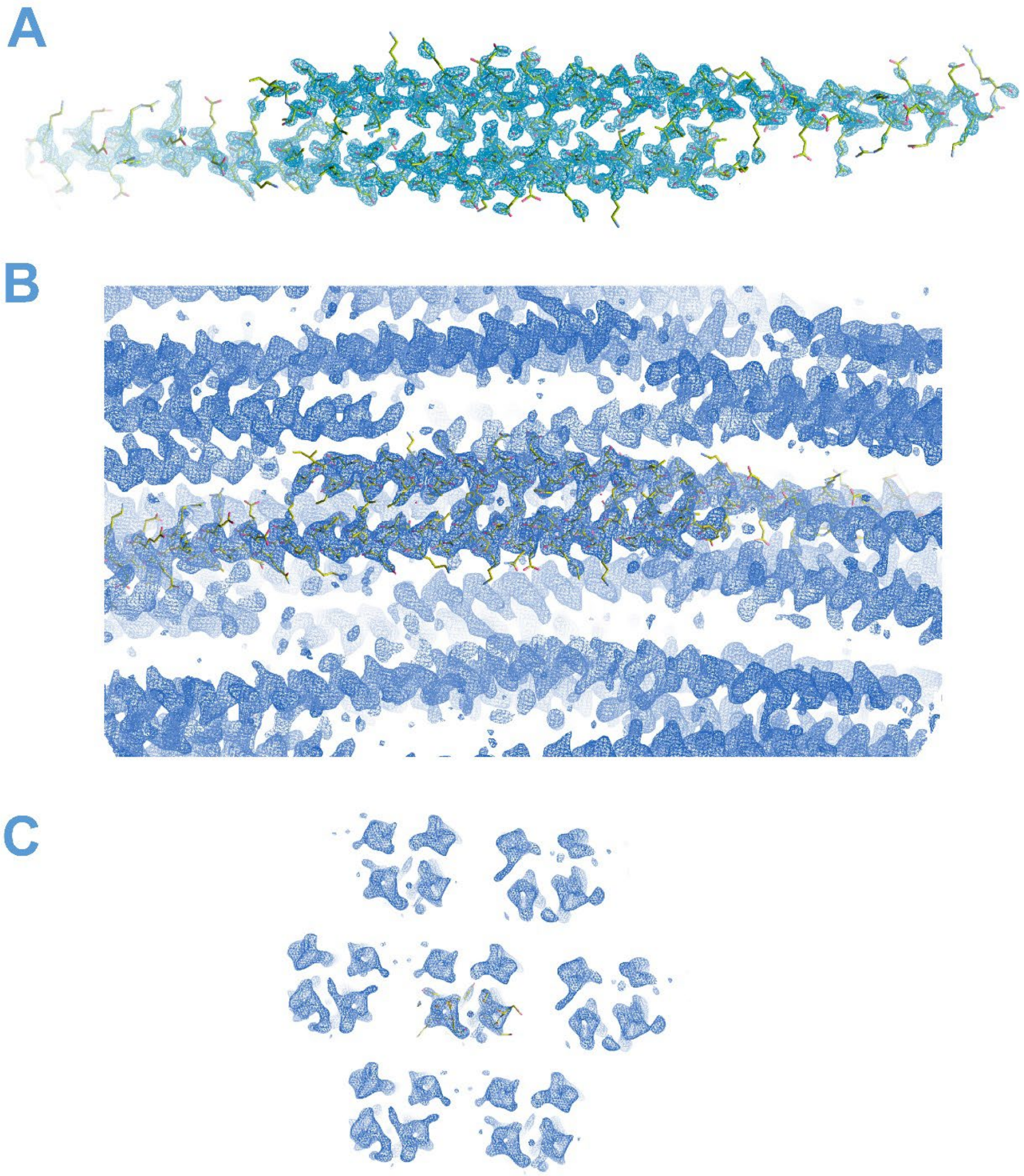

***Supplementary Figure 12 – Electron density of the Myo6 875-940 crystal structure.***

(A) Electron density corresponding to the 875-940 dimer (2Fo-Fc map contoured at 1.0 RMSD).
(B) View of the 2-fold axis that defines the antiparallel dimer.
(C) View of the 6-fold axis. Although a high number of copies can be seen in the asymmetric unit, the 875-937 helix makes extensive contacts over 11.5 turns with another helix. The rest of the crystal contacts made by the helix are negligible in comparison (Sup. Movie 2). Images prepared with Coot 0.9.8.1[7].

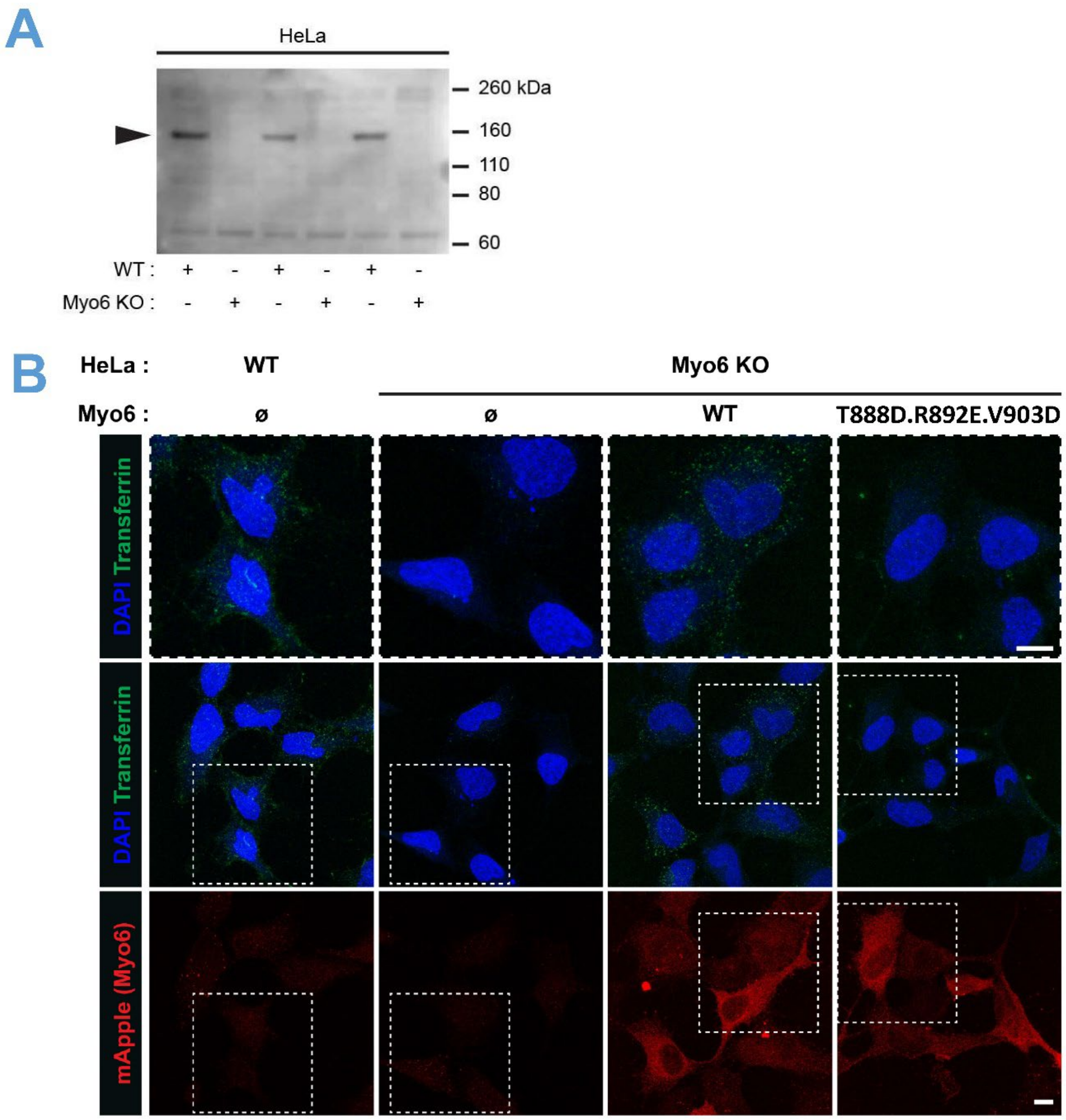


***__Supplementary Figure 13 – Endocytosis is impaired if Myo6 cannot dimerize through its proximal region__***

**(A)** Confirmation of Myo6 null HeLa cells by western blot.

**(B)** Endocytic uptake of fluorescently-labeled transferrin by WT and Myo6 null HeLa cells. Internalized transferrin appears in green, HeLa cell nuclei (DAPI) in blue, and expression of $^{\text{mApple}}$Myo6 constructs in red. Representative maximum-intensity projection images demonstrate a marked reduction in internalized transferrin in Myo6 mutant HeLa cells compared to wild-type cells. Forced expression of FLMyo6 (WT), but not of FLMyo6 (T888D.R892E.V903D, triple mutant), restores the ability of Myo6 null HeLa cells to uptake transferrin. Scale bar: 10 µm.

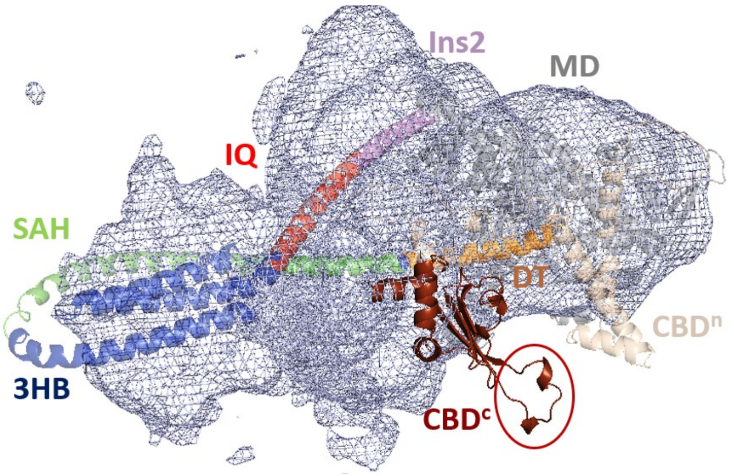


***Supplementary Figure 14 – Alphafold[8,9] predicts a back-folded FLMyo6 with an exposed CBD$^c$***

Alphafold modelisation of FLMyo6 (UNIPROT: Q9UM54) was manually docked within our 3D reconstruction of Jo-Myo6-In (grey mesh). CBD$^c$ is found outside of the electron density, with residues D1157, Y1159, D1161, Q1163 highly exposed (red circle), inconsistently with our experimental data.

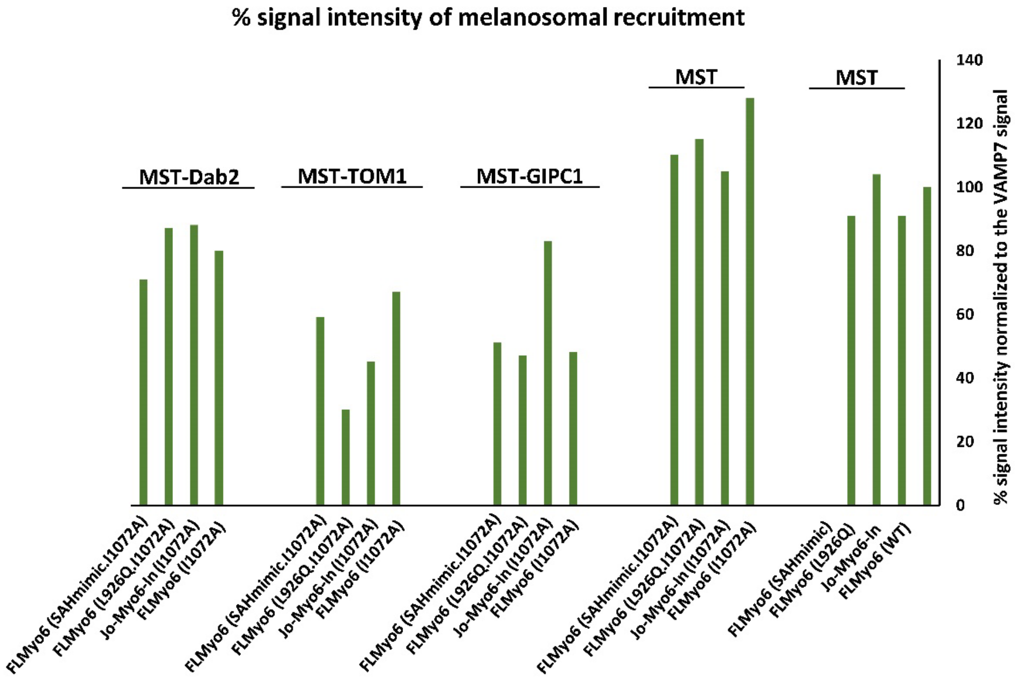


***Supplementary Figure 15 – Fluorescence of MST-partners localized on melanosomes, normalized to the VAMP7 signal.***

Quantification of the fluorescence signal of $^{mCherry}$MST-partners on melanosomes in MNT-1 cells expressing different Myo6 constructs. Global intensities were calculated by using the macro described in Methods, normalized to the $^{iRFP}$VAMP7 signal (total melanosomal surface area).

| | |
|---|---|
| **Space group** | $P6_522$ |
| **Cell constants**<br>**a, b, c, α, β, γ** | 29.39 Å, 29.39 Å, 295.65 Å<br>90.00°, 90.00°, 120.00° |
| **Resolution** | 49.27 Å - 2.22 Å |
| **Data completeness in resolution range** | 66.9 % |
| **$R_{merge}$** | 0.17 |
| **I/s(I)** | 1.02 at 2.22 Å |
| **R, Rfree** | 0.304, 0.343 |
| **Wilson B-factor** | 39.2 $Å^2$ |
| **Average B, all atoms** | 55.0 $Å^2$ |
| **Total number of atoms** | 1051 |
| **Unique reflections** | 2977 |
| **Rfree test set** | 301 reflections (10.1%) |
| **Ramachandran outliers** | 0 |
| **Sidechain outliers** | 8.8% |
| **Solvent content*** | 45.1% |

***Supplementary Table 1 – Crystallographic structure of the Myo6 dimerization domain – data and refinement statistics.***

* Solvent content estimated by *Matthews_coef*[10]

| Construct | IMAC | SEC |
|---|---|---|
| Ins2/IQ/3HB* | 20 mM Tris pH 7.5; 150 mM NaCl; 1 mM DTT (+ 4 and 300 mM Imidazole for binding/washing and elution, respectively) | 20 mM HEPES, pH 7.5; 50 mM NaCl; 2.5mM $MgCl_2$; 1 mM TCEP |
| $^{YFP}$CBD (WT and mutant) | 20 mM Tris pH 7.5; 150 mM KCl; 40 mM Imidazole pH 7.5; 5% Glycerol; 1mM DTT (+300 mM NaCl in the washing step; and 100 mM KCl; 300 mM imidazole; 1 mM TCEP for elution) | 20 mM Tris pH 7.5; 100 mM KCl; 1 mM TCEP |
| GIPC1 | 20 mM HEPES pH 7.5; 150 mM NaCl; 1 mM DTT (+4 or 200 mM Imidazole for binding/washing and elution, respectively) | 50 mM HEPES pH 7.5; 150 mM or 50 mM NaCl; 1 mM DTT |
| TOM1 | 20 mM Tris pH 7.5; 300 mM NaCl; 1 mM DTT (+4 or 300 mM Imidazole for binding/washing and elution, respectively) | 20 mM HEPES pH 7.5; 150 mM NaCl; 1 mM DTT |
| Dab2 | 20 mM Tris pH 8.5; 300 mM NaCl; 1 mM DTT (+4 or 200 mM Imidazole for binding/washing and elution, respectively) | 20 mM Tris pH 8.5; 150 mM NaCl; 1 mM DTT |
| Myo6 875-940, 875-955**, 880-940, 888-940** and 834-955 | 20 mM Tris, pH7.5; 150 mM NaCl (+4 or 200 mM Imidazole for binding/washing and elution, respectively) | 10 mM Tris, pH7.5 ; 50mM NaCl |
| Myo6 875-940 triple mutant** | 20 mM Tris, pH7.5; 600 mM NaCl (+4 or 200 mM Imidazole for binding/washing and elution, respectively) | 10 mM Tris, pH7.5 ; 300mM NaCl |

***Supplementary Table 2 – Purification buffers used for constructs expressed in E. Coli***

* presence of 2 CaM bound was confirmed by SEC-MALS (not shown).
** Ion exchange in MonoQ 5/50 GL prior to SEC.
*** Ion exchange in MonoS 5/50 GL prior to SEC.

(For constructs including a rTEV cleavage site, homemade rTEV was added 1:50 m/m, and a second passage through IMAC was performed before SEC)

### Supplementary Movies

***Supplementary Movie 1***

3D view of the Myo6 off-state docked in the negative staining reconstruction (from Fig 3A).

***Supplementary Movie 2***

Representation of the crystal packing of the dimerization Myo6 fragment 875-937 as calculated with PyMol. Each crystallographic dimer is represented in a different color.

### Supplementary Material and Methods

#### *Cloning*, expression and purification from SF9/baculovirus system.

The full-length wild-type Myo6 (FLMyo6) was generated using human Myo6 cDNA splice form without the large insertion (Q9UM54-2 in UNIPROT). The small insert was removed through sub cloning to obtain a FLMyo6 construct without any spliced insert (corresponding to isoform Q9UM54-5). The FLMyo6 (no inserts) construct was then used in all *in vitro* experiments requiring a full-length construct except for the Anti-His pull-down experiment, for which the small insert isoform was used.
The deafness mutant (L926Q) and triple mutant (T888D.Q892E.V903D - R892 in mouse corresponds to Q892 in human, see Sup Fig. 10B) in the anti-parallel dimerization region were produced from FLMyo6 with no inserts by reverse PCR. As previously described[11], a mutant FLMyo6 (SAHmimic) was made where the residues from Glu922 to Glu935 (EAERLRRIQEEMEK) were replaced with alternate acidic and basic residues (EERKRREEEERKKREEE) to match the (i, i+4) phasing observed in the predicted Myo6 SAH domain.
For microscale thermophoresis and ATPase assays, previously described constructs were used: MD (1-789)[12], $MD^{Ins2}$ (2-816)[3], $MD^{Ins2/IQ/3HB}$(1-917)[13]. The Myo6 zippered dimer was created by truncation at R991 followed by a leucine zipper (PDB: GCN4[14]) as previously described[15].
For the bundle unfolding experiments, a monomeric "cys-lite" construct was made by C-terminal truncation at amino acid Q919 and introduction of C321S, C362S, and C611A. To this construct, either a T845C mutation alone, or the combination of T845C and A880C mutations was introduced for rhodamine labeling, as previously described[13]. A dimeric "cys-lite" construct was made by introduction of the C321S, C362S, C611A mutations in the Myo6 zippered dimer[15]. Into this construct, either a T845C mutation alone, or the combination of T845C and A880C mutations were introduced for rhodamine labeling, as previously described[13], with or without the addition of a deafness-causing mutation (L926Q).
Each of these constructs had a Flag tag (GDYKDDDDK) at its N-terminal end to facilitate purification. Expression in baculovirus system and purification were performed as previously described[16].
Jo and In-Flag sequences[4] were synthesized (Eurofins genomics) and fused to Myo6 N-terminus (linker Gly-Ser) and C-terminus (linker Gly), in pVL1392 for expression in Sf9 cells. Purification was achieved using the published protocol for FLMyo6[16] except that, for EM studies, purification was performed by replacing ATP with $ADP.VO_4$ in the lysis buffer. For increased purity, a SEC step was performed using a Superdex 200 Increase column (Cytiva) developed in 10 mM Hepes; 80 mM NaCl; 5mM $NaN_3$; 1 mM $MgCl_2$; 0.1 mM TCEP; 0.1 mM ADP; 0.2 mM $VO_4$; 0.1 mM EGTA; pH 7.5.

#### *Constructs cloning, expression and purification from Escherichia coli*

##### *Cloning Myo6 constructs*

Ins2/IQ/3HB was generated using our human FLMyo6 NI construct (see previous section), DNA sequence encoding for aa783-917 was transferred into pPROX-HTB plasmid containing in N-terminus 6XHis tag and a TEV cleavage sequence (coding for ENLYFQG).
Myo6 $^{YFP}$CBD was generated through several round of subclonings from cDNA mouse Myo6 (E9Q3L1 in UNIPROT). Myo6 was incorporated in pET14 plasmid containing in N-terminus 6XHis-tag, yellow fluorescent protein (YFP) and a TEV cleavage sequence. Finally, Myo6 was truncated in N-terminus at position corresponding to aa M1032 through reverse PCR. The $^{YFP}$CBD (D1157V.Y1159D.D1161R.

Q1163V) mutant was generated through point mutations addition using reverse PCR on the $^{YFP}$CBD (WT) construct.

##### *Cloning partner constructs*

In order to avoid partner (GIPC1, TOM1 and Dab2) degradation and auto-inhibition as previously reported[17,18], we used truncations containing the published Myo6-binding domains[18–21] instead of FL constructs.

For microscale thermophoresis assays and his-pull-down assays, $^{His}$GIPC1 construct was generated using cDNA full length mouse GIPC1 (UNIPROT Q9Z0G0). GIPC1 was incorporated in pProEX-HTb plasmid containing in N-terminus 6XHis-tag and a TEV cleavage sequence. Finally, GIPC1 was truncated in N-terminus at position corresponding to aa D255 through reverse PCR in order to keep only the GH2 domain which is sufficient for mediated interaction with Myo6[18].

For ATPase assays, we used GIPC1 in fusion with the mNeonGreen tag: GIPC1 DNA sequence encoding for residues 238-end was incorporated in the pET28 plasmid containing in the N-terminus 6XHis-tag and a mNeonGreen tag using homemade Gibson Assembly mix[22].

For microscale thermophoresis and his-pull-down assays, a non-fluorescent $^{His}$TOM1 construct was generated using cDNA full length human TOM1 (UNIPROT O60784-1). TOM1 207-end was incorporated into pET14 in-frame with an N-terminus 6XHis-tag and a TEV cleavage sequence using homemade Gibson Assembly mix[22].

For ATPase assays, we used the TOM1 436-461 peptide described as the minimal sequence required for TOM1 binding to Myo6[21]. DNA sequence coding for aa436-461 was incorporated in pET14 plasmid containing in N-terminus 6XHis-tag, yellow fluorescent protein (YFP) and a TEV cleavage sequence.

Dab2 His-650-end (tDab2[17]) was a kind gift of Christopher Toseland.

To identify the minimal sequence involved in proximal dimerization, several Myo6 truncations were generated from mouse Myo6 cDNA (UNIPROT: E9Q3L1). Constructs encoding for aa 875-940 and 875-955 were cloned with a N-terminal 6xHis-tag into pET14 plasmid. The construct encoding for aa 834-955 was cloned into pET14 with a N-terminal 6xHis-tag followed with a Thrombin cleavage site (coding for LVPRGSH). Constructs encoding for aa 880-940 and 888-940 were cloned with a C-terminal 6xHis-tag into pET14 by PCR and blunt-end ligation. The 912-end construct was cloned into pProEX-HTb with a N-terminal 6xHis-tag through several rounds of sub cloning. For crystallization assays, the construct encoding for aa 875-940 was generated with a N-terminal 6xHis-tag and a TEV cleavage sequence into pProEX-HTb using homemade Gibson Assembly mix[22]. The point mutations T888D, R892E, V903D were added in the backbone encoding for his-rTEV-875-940, all by Quickchange[23].

##### *Protein expression and purification*

Constructs were expressed in E.coli BL21 (DE3) cells (NEB). Cells were grown in 2xYT media until $OD_{560}$~0.8, expression was then induced by addition of 200 µM isopropyl β-D-1-thiogalactopyranoside (except for Dab2 expression, where 1 mM isopropyl β-D-1-thiogalactopyranoside was used). Cells were lysed by sonication. For purification, the lysate soluble fraction was loaded on an IMAC column (cOmplete 5mL, Roche for all constructs except $^{YFP}$CBD (WT and mutant) for which HisTrap-FFcrude 5mL, Cytiva was used instead), and proteins were eluted with 200 mM or 300 mM Imidazole. Purest fractions were identified by SDS-PAGE. If needed, pooled fractions were concentrated using Vivaspin concentrators (Sartorius) up to ~5 mL. Concentrated samples were injected in Superdex 200 or 75 16/600 columns (Cytiva) depending on the molecular weight of the target protein. Purest fractions and the final sample were concentrated by ultrafiltration, and protein concentration was determined using

Nanodrop 2000 (ThermoScientist). The final sample containing concentrated protein was flash-frozen in liquid nitrogen and stored at -80°C.

For proteins containing a TEV cleavable His-tag, prior to the gel filtration, His-tag was removed by incubation with homemade rTEV protease overnight in a 1/50 mass ratio. The incubate was passed through the cOmplete His-Tag Purification Column again to remove rTEV and the uncleaved fraction, then concentrated and loaded in a Superdex 75-16/60 gel filtration column.

Purification buffers are detailed in Sup Table 2.

***Constructs cloning for expression in MNT-1 cells***

For expression in MNT-1, FLMyo6 was generated from cDNA of full length human Myo6, no inserts isoform (UNIPROT: Q9UM54-5) with shRNA resistance. DNA was transferred to the pEGFP-C1 vector via a XbaI restriction enzyme site. SAHmimic mutations (Glu922 to Glu935 (EAERLRRIQEEMEK) replaced with alternate acidic and basic residues (EERKRREEEERKKREE) were introduced by reverse PCR. The L926Q mutation was introduced using Quickchange[23]. Transfer of Jo-Myo6-In from baculovirus vector to P-EGFP-C1 was ordered from GenScript. Myo6 CBD was generated by transferring DNA encoding G1037-end from human Myo6, no inserts isoform (UNIPROT: Q9UM54-5) into pEGFP-C1 plasmid using the XbaI restriction enzyme site.

I1072A was introduced in previously cloned constructs (see above) using reverse PCR.

Mouse GIPC1 (239-end), human TOM1 (299-end as described in[19]) and human Dab2 (650-end) were transferred in a modified pmCherry-C1 plasmid containing in N-terminus a melanosome-targeting tag (MST tag, aa 1-139 from Mouse MREG – UNIPROT: Q6NVG5) as described in [24]. The MST tag and mCherry are separated by a GGSGGTGG linker. In the $^{mCherry}$MST-partners constructs, mCherry and GIPC1, TOM1 or Dab2 sequences are separated by the polylinker multiple cloning site SGLRSRAQASNSLTSK.

### References


1. Manalastas-Cantos, K. *et al.* ATSAS 3.0: expanded functionality and new tools for small-angle scattering data analysis. *J Appl Cryst* **54**, 343–355 (2021).
2. Berggren, K. *et al.* Background-free, high sensitivity staining of proteins in one- and two-dimensional sodium dodecyl sulfate-polyacrylamide gels using a luminescent ruthenium complex. *Electrophoresis* **21**, 2509–2521 (2000).
3. Ménétrey, J. *et al.* The structure of the myosin VI motor reveals the mechanism of directionality reversal. *Nature* **435**, 779–785 (2005).
4. Bonnet, J. *et al.* Autocatalytic association of proteins by covalent bond formation: a Bio Molecular Welding toolbox derived from a bacterial adhesin. *Sci Rep* **7**, 43564 (2017).
5. Barnes, C. A. *et al.* Remarkable Rigidity of the Single α-Helical Domain of Myosin-VI As Revealed by NMR Spectroscopy. *J Am Chem Soc* **141**, 9004–9017 (2019).
6. Kim, H., Hsin, J., Liu, Y., Selvin, P. R. & Schulten, K. Formation of salt bridges mediates internal dimerization of myosin VI medial tail domain. *Structure* **18**, 1443–1449 (2010).
7. Emsley, P. & Cowtan, K. Coot: model-building tools for molecular graphics. *Acta Crystallogr D Biol Crystallogr* **60**, 2126–2132 (2004).
8. Jumper, J. *et al.* Highly accurate protein structure prediction with AlphaFold. *Nature* **596**, 583–589 (2021).
9. Varadi, M. *et al.* AlphaFold Protein Structure Database: massively expanding the structural coverage of protein-sequence space with high-accuracy models. *Nucleic Acids Res* **50**, D439–D444 (2022).
10. Matthews, B. W. Solvent content of protein crystals. *J Mol Biol* **33**, 491–497 (1968).
11. Mukherjea, M. *et al.* Myosin VI must dimerize and deploy its unusual lever arm in order to perform its cellular roles. *Cell Rep* **8**, 1522–1532 (2014).
12. Ménétrey, J., Llinas, P., Mukherjea, M., Sweeney, H. L. & Houdusse, A. The structural basis for the large powerstroke of myosin VI. *Cell* **131**, 300–308 (2007).
13. Mukherjea, M. *et al.* Myosin VI dimerization triggers an unfolding of a three-helix bundle in order to extend its reach. *Mol Cell* **35**, 305–315 (2009).
14. Lumb, K. J., Carr, C. M. & Kim, P. S. Subdomain folding of the coiled coil leucine zipper from the bZIP transcriptional activator GCN4. *Biochemistry* **33**, 7361–7367 (1994).
15. De La Cruz, E. M., Ostap, E. M. & Sweeney, H. L. Kinetic mechanism and regulation of myosin VI. *J Biol Chem* **276**, 32373–32381 (2001).
16. Sweeney, H. L. *et al.* Kinetic tuning of myosin via a flexible loop adjacent to the nucleotide binding pocket. *J Biol Chem* **273**, 6262–6270 (1998).
17. Fili, N. *et al.* Competition between two high- and low-affinity protein-binding sites in myosin VI controls its cellular function. *J Biol Chem* **295**, 337–347 (2020).
18. Shang, G. *et al.* Structure analyses reveal a regulated oligomerization mechanism of the PlexinD1/GIPC/myosin VI complex. *Elife* **6**, e27322 (2017).
19. Tumbarello, D. A. *et al.* Autophagy receptors link myosin VI to autophagosomes to mediate Tom1-dependent autophagosome maturation and fusion with the lysosome. *Nat Cell Biol* **14**, 1024–1035 (2012).
20. Yu, C. *et al.* Myosin VI undergoes cargo-mediated dimerization. *Cell* **138**, 537–548 (2009).
21. Hu, S. *et al.* Structure of Myosin VI/Tom1 complex reveals a cargo recognition mode of Myosin VI for tethering. *Nat Commun* **10**, 3459 (2019).
22. Gibson, D. G. *et al.* Enzymatic assembly of DNA molecules up to several hundred kilobases. *Nat Methods* **6**, 343–345 (2009).
23. Liu, H. & Naismith, J. H. An efficient one-step site-directed deletion, insertion, single and multiple-site plasmid mutagenesis protocol. *BMC Biotechnol* **8**, 91 (2008).
24. Ishida, M., Arai, S. P., Ohbayashi, N. & Fukuda, M. The GTPase-deficient Rab27A(Q78L) mutant inhibits melanosome transport in melanocytes through trapping of Rab27A effector protein Slac2-a/melanophilin in their cytosol: development of a novel melanosome-targetinG tag. *J Biol Chem* **289**, 11059–11067 (2014).
